# Decoding the molecular logic of rapidly evolving ZAD zinc-finger proteins in *Drosophila*

**DOI:** 10.1101/2025.05.21.655257

**Authors:** Raku Saito, Yusuke Umemura, Shiho Makino, Takashi Fukaya

**Affiliations:** Laboratory of Transcription Dynamics, Research Center for Biological Visualization, Institute for Quantitative Biosciences, The University of Tokyo, Bunkyo-ku, Tokyo 113-0032, Japan; Department of Life Sciences, Graduate School of Arts and Sciences, The University of Tokyo, Bunkyo-ku, Tokyo 113-0032, Japan

## Abstract

The zinc-finger associated domain (ZAD)-containing C2H2 zinc-finger proteins (ZAD-ZnFs) represent the most abundant class of transcription factors that emerged during insect evolution, yet their molecular diversity and biological functions remain largely unclear. Here, we established a systematic CRISPR-based protein-tagging approach that enables direct, unambiguous comparison of nuclear localization and genome-wide binding profiles of endogenous ZAD-ZnFs in developing *Drosophila* embryos. Evidence is provided that a subset of ZAD-ZnFs forms nuclear condensates through the stacking of the N-terminal ZAD dimerization surface. Disruption of condensation activity leads to misregulation of genome-wide binding profiles and lethality, underscoring its functional and physiological significance in development. Importantly, integrative ChIP-seq and Micro-C data analyses reveal that many ZAD-ZnFs colocalize with core insulator proteins such as CTCF and CP190 to strengthen the formation of topological boundaries. We suggest that the diverse molecular functions of ZAD-ZnFs have evolutionally arisen from their ancestral role as insulator-binding proteins.

## Introduction

The zinc-finger associated domain (ZAD)-containing C2H2 zinc-finger proteins (ZAD-ZnFs) represent the most abundant class of transcription factors that has emerged during insect evolution.^1^ ZAD-ZnFs consist of an N-terminal ZAD and arrays of C2H2 zinc-finger domains at the C-terminus, flanked by flexible linker regions. Similar to the POZ/BTB and SCAN domains in mammals and other species, the N-terminal ZAD functions as a protein dimerization scaffold through the stacking of the hydrophobic surface.^2–6^ More than 90 ZAD-ZnF genes are present in the *Drosophila melanogaster* genome, but the number of ZAD-ZnF genes greatly varies across related *Drosophila* species.^7^ Despite such rapid evolutionary dynamics,^1,7,8^ ZAD-ZnF genes often encode proteins with essential biological functions. For example, ZAD-ZnF genes *Molting Defective*, *Ouija Board*, and *Séance* encode transcriptional activators responsible for the expression of essential ecdysone biosynthesis genes during larval development.^9,10^ On the other hand, evolutionally young ZAD-ZnF genes *Oddjob* (*Odj*), *Nicknack* (*Nnk*), and *Identity crisis* (*Idc*) encode DNA-binding proteins that associate with heterochromatic regions enriched with the repressive H3K9me3 histone mark.^7,11,12^ The ZAD-ZnFs Kipferl (Kipf) and Trailblazer (Trabl) were recently reported to function as positive regulators of the Piwi-piRNA-mediated transposon silencing pathway in reproductive tissues.^13,14^ In addition, the ZAD-ZnFs M1BP, Pita, ZIPIC, and Zw5 are thought to help organize higher-order genome topology through the association with insulator elements.^15–17^ As an exceptional case, a ZAD-ZnF gene *weckle* (*wek*) is suggested to encode a membrane-anchored cytoplasmic protein that acts as an adaptor protein in the Toll pathway during the establishment of the dorsal-ventral axis in early embryos.^18^ Overall, these findings are consistent with the idea that individual ZAD-ZnFs have functionally diverged to control widespread biological processes during the evolution of *Drosophila* species under natural environments. However, the evolutionary origin and common molecular features of ZAD-ZnFs remain uncertain.

As exemplified above, extensive efforts have been made to elucidate the biological roles and molecular properties of individual ZAD-ZnF genes over decades. However, we are still far from a comprehensive understanding of their *in vivo* functions, since the systematic characterization of endogenous ZAD-ZnFs in a unified experimental setup is critically lacking. To overcome this hurdle, we have established a unified protein-tagging system that enables comprehensive, direct comparison of nuclear localization, genome-wide binding profiles, and *in vivo* functionalities of endogenous ZAD-ZnFs in developing *Drosophila* embryos. CRISPR/Cas9-mediated insertion of a DNA cassette encoding a GFP-3xFLAG tag immediately upstream of the stop codon permits super-resolution live-imaging and ChIP-seq analyses of individual ZAD-ZnFs by minimizing potential experimental noise arising from the fixation protocol for immunostaining^19^ and the variability of antibody quality for ChIP experiments.^20,21^ Using our unique experimental framework, we identified a distinct subset of ZAD-ZnFs that forms nuclear condensates via the N-terminal ZAD dimerization surface. Disruption of condensation activity leads to misregulation of genome-wide binding profiles and lethality, underscoring its functional and physiological significance in development. We also show that a class of ZAD-ZnFs exhibiting uniform nuclear localization does not rely on the N-terminal ZAD to exert their biological functions, highlighting rapid functional and molecular divergence within this family. Importantly, integrative analysis of Micro-C data along with ChIP-seq datasets reveals that many ZAD-ZnFs colocalize with well-characterized core insulator proteins such as CTCF, Su(Hw), BEAF-32, and CP190 to strengthen the formation of topological boundaries, suggesting that insulator-binding activity is a common ancestral feature of ZAD-ZnFs. We suggest that ZAD-ZnFs constitute a novel class of genome organizers with diverse molecular properties, likely shaped by rapid functional divergence during insect evolution.

## Result

### Establishment of CRISPR/Cas9-mediated protein-tagging system

To systematically visualize the localization patterns of endogenous proteins in living *Drosophila*, a sequence cassette encoding GFP-3xFLAG or mCherry-3xFLAG tag has been individually inserted immediately upstream of the stop codon of target genes (Figure S1A). Resulting genome-edited strains were then crossed with a Cre-expressing strain to remove the *3xP3-dsRed* selection marker from the targeted locus. Insertion of the desired sequence cassette has been verified by PCR analysis of purified genomic DNA for all the genome-edited strains produced in this study. As a test example, we initially targeted the well-characterized *vasa* gene, which is essential for the development of ovarian tissues through the formation of a perinuclear compartment called nuage in nurse cells.^22,23^ A western blotting signal against an anti-FLAG antibody was specifically seen upon genome editing (Figure S1B). In addition, nuage formation was reproducibly observed by live-imaging analysis of Vasa-mCherry-3xFLAG fusion protein in dissected ovaries (Figure S1C). Importantly, the embryo hatching rate was essentially unchanged even after genome-editing of the *vasa* gene (Figure S1D), supporting the idea that our CRISPR/Cas9-mediated tagging approach enables live visualization of protein products of endogenous key developmental genes in a non-interfering manner.

As another test example, we next focused on the well-characterized DNA-binding protein GAGA factor (GAF), which is encoded by the *Trithorax-like* (*Trl*) gene in *Drosophila*.^24,25^ A number of imaging studies reported that GAF protein forms a cluster or condensate within a nucleus in developing embryos.^e.g., 26–29^ A sequence cassette encoding either GFP-3xFLAG or mCherry-3xFLAG tag has been individually inserted immediately upstream of the stop codon at the endogenous *Trl* locus. Resulting genome-edited strains were both homozygous viable, suggesting that the inserted GFP-3xFLAG or mCherry-3xFLAG tag does not impede either transcription or translation. As in the case of *vasa*, a clear western blotting signal against an anti-FLAG antibody was specifically detected upon genome-editing (Figure S1E). Consistent with previous literature,^27–29^ both GAF-GFP-3xFLAG and GAF-mCherry-3xFLAG formed a cluster within a nucleus in developing embryos at nuclear cycle 14 (nc14) (Figure S1F), indicating that the unique localization patterns of endogenous protein products are not affected by the choice of fluorescent proteins. Importantly, condensate-like localization was not seen when GFP-3xFLAG tag alone was expressed in conjunction with an SV40 nuclear localization signal (Figure S1G), ruling out the possibility that the inserted GFP-3xFLAG tag itself drives condensate assembly in a non-specific manner.

### Visualization of endogenous insulator proteins in early embryos

Having established our CRISPR/Cas9-mediated protein-tagging strategy, we first sought to explore the localization patterns of so-called insulator proteins, a class of nuclear proteins that plays a central role in organizing the 3D genome structure in *Drosophila* (reviewed in Bortle and Corces^30^). In this study, we particularly focused on six well-characterized insulator proteins, namely Su(Hw), CTCF, BEAF-32, CP190, L(3)mbt, and Fs(1)h-L.^31–36^ As in the case of GAF, a GFP-3xFLAG tag has been inserted immediately upstream of the stop codon of individual target genes. All the resulting genome-edited strains were homozygous viable, suggesting that the GFP-3xFLAG tag does not impede the functionality of these targeted genes. Successful tagging was confirmed by western blot analysis using anti-FLAG antibody (Figure S2A). Previous studies suggested that insulator proteins form condensates called “insulator bodies” within the nucleus of dissected imaginal discs or in cultured cells.^e.g., 34,37–42^ However, it remains uncertain whether endogenous insulator proteins actually form “insulator bodies” in living *Drosophila*, especially given a recent finding that the traditional fixation protocol for immunostaining can significantly alter the appearance of protein condensates.^19^ Intriguingly, Airyscan super-resolution live-imaging analysis of the newly produced genome-edited strains revealed that each insulator protein exhibits distinct patterns of nuclear localization in early embryos at the same developmental stage (∼15 min after entry into nc14) (Figure 1A). For example, Su(Hw) has long been postulated as a major constituent of “insulator bodies”,^37–39,42^ but its localization pattern was more uniform than those of other insulator proteins such as CP190 and Fs(1)h-L (Figure 1A). In addition, unlike these proteins, CTCF and L(3)mbt formed only a few bright foci per nucleus, while BEAF-32 was more uniformly distributed in the cytoplasm and nucleus without assembling clear nuclear foci (Figure 1A). Taken these results together, it is not likely that all the tested insulator proteins coalesce to form common “insulator bodies” to regulate 3D genome organization in developing embryos. This view is consistent with a recent ChIP-seq study reporting distinct distribution profiles of insulator proteins in nc14 embryos.^43^ We will use our newly produced genome-edited insulator strains for subsequent ChIP-seq analysis, which will be discussed in the later part of this study (Figures 6 and 7).

**Figure 1.**
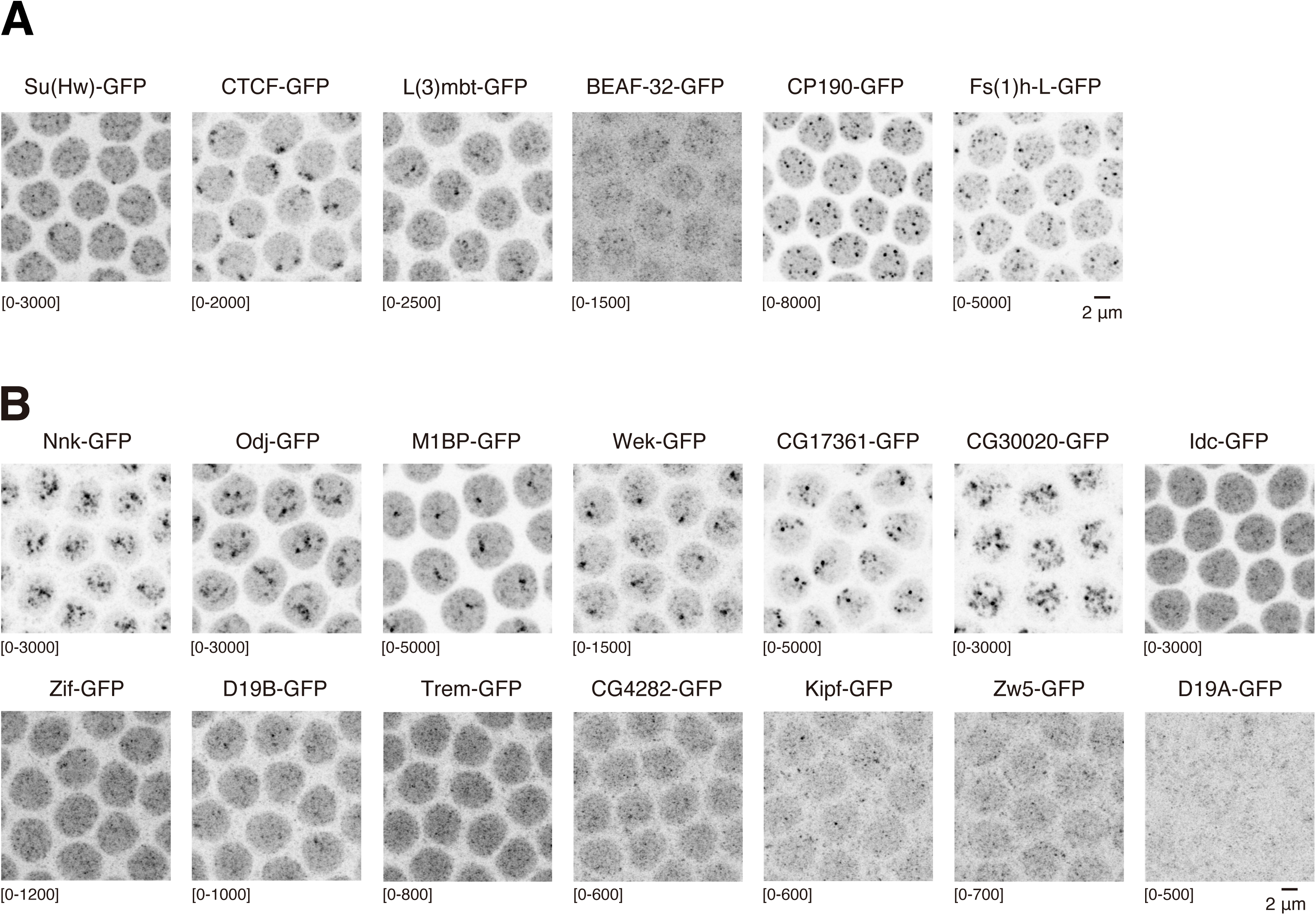
Live-visualization of endogenous insulator proteins and ZAD-ZnFs in early embryos. (A) Airyscan imaging of GFP-3xFLAG-tagged endogenous insulator proteins in living embryos. Images were taken ∼15 min after entry into nc14. Embryos from corresponding homozygous genome-edited strains were used for the analysis. The maximum intensity projected images are shown. (B) Airyscan imaging of GFP-3xFLAG-tagged endogenous ZAD-ZnFs in living embryos. Images were taken ∼15 min after entry into nc14. Embryos from corresponding homozygous genome-edited strains were used for the analysis. The maximum intensity projected images are shown. See Figure S1 and S2.

### Systematic visualization of endogenous ZAD-ZnFs in early embryos

To systematically characterize the localization patterns of ZAD-ZnFs in *Drosophila*, we targeted 14 ZAD-ZnF genes expressed in early embryos for CRISPR/Cas9-mediated insertion of the GFP-3xFLAG tag at the C-terminal end (Figure S1A). Successful tagging was confirmed by western blot analysis using anti-FLAG antibody (Figure S2B). While all the ZAD-ZnFs share a common structural arrangement (*i.e.*, an N-terminal ZAD followed by a flexible unstructured linker and an array of C-terminal C2H2 zinc-finger domains), their nuclear localization was found to be very different (Figure 1B). For example, Nnk, Odj, M1BP, Wek, CG17361, and CG30020 reproducibly formed a cluster or condensate within a nucleus, while others (Idc, Zif, D19B, Trem, CG4282, Kipf, Zw5, and D19A) were more uniformly distributed in nc14 embryos. Among these, M1BP and Zw5 are well characterized as insulator proteins.^16,17,44,45^ We noticed that the localization patterns of M1BP and Zw5 are distinct from those of non-ZAD-ZnF insulator proteins such as CP190, Fs(1)h-L, and others at the same developmental stage (Figure 1A), further reinforcing the idea that hypothetical “insulator bodies” do not represent co-condensation of all insulator proteins at the same site to shape 3D genome topology. Another striking finding here is the nuclear localization of Wek in nc14 embryos (Figure 1B), since a previous study reported that Wek specifically localizes to the plasma membrane and acts as an adaptor of the Toll pathway to control dorsal-ventral patterning at this developmental stage.^18^ Our subsequent ChIP-seq analysis further supports the idea that Wek acts as a nuclear DNA-binding protein that associates with insulator elements (Figures 6 and 7), challenging the traditional view that Wek exerts its function as a cytoplasmic membrane-anchored adaptor protein of the Toll pathway in early embryos.

### ZAD is required for the *in vivo* function of Nnk and CG30020/BroN

To elucidate the mechanism underlying the clustering of ZAD-ZnFs and its biological significance, we decided to focus on two ZAD-ZnFs, Nnk and CG30020, as they form clear clusters within a nucleus (Figure 1B). Previous biochemical and X-ray crystallography studies reported that the N-terminal ZAD mediates homotypic protein-protein interactions through the stacking of a hydrophobic surface (Figure S3A and B: left).^2,5,6^ We initially tested whether this dimerization activity of ZAD could be reliably predicted using the newly developed AlphaFold3 algorithm.^46^ It turned out that the previously reported X-ray crystal structure of ZAD dimers was nicely recapitulated, including the positioning of the flexible regions and the Zn²⁺ ion coordination sites (Figure S3A and B: right). Then, the AlphaFold3 algorithm was applied for the analysis of Nnk. It was predicted that its ZAD also forms a dimer with high confidence (Figure 2A and B), in a way very similar to the experimentally verified Grau and M1BP ZAD dimers (Figure S3A-B). To examine the biological role of ZAD, Nnk WT and ΔZAD mutant were individually expressed as a GFP-3xFLAG fusion protein under the control of the native *Nnk* promoter (Figure 2C). Intriguingly, cluster formation of Nnk was completely abolished upon the loss of the N-terminal ZAD (Figure 2D), indicating that the dimerization activity of ZAD is required for the assembly of the Nnk cluster in early embryos. We then analyzed CG30020 following the same experimental strategy (Figure 2E). As in the case of Nnk, AlphaFold3 predicted that the CG30020 ZAD also serves as a dimerization scaffold (Figure 2F). When CG30020 WT and ΔZAD mutant were individually expressed as a GFP-3xFLAG fusion protein under the control of the native *CG30020* promoter (Figure 2G), only the WT construct was able to assemble a cluster within a nucleus (Figure 2H). Based on molecular similarities between CG30020 and Nnk, we named *CG30020* as *Brother of Nicknack*, or *BroN* (Figure 2E).

**Figure 2.**
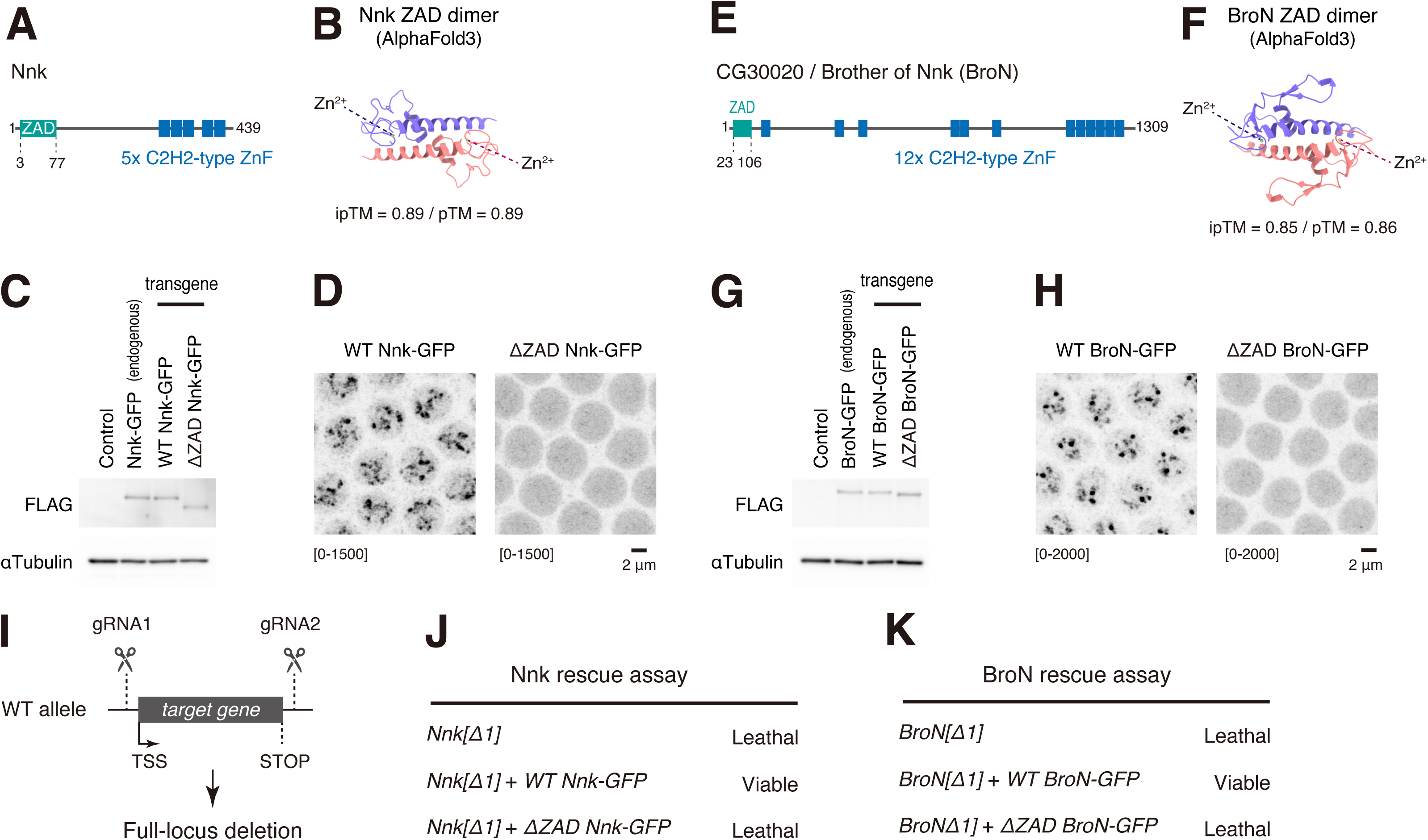
The N-terminal ZAD is essential for Nnk and BroN functions. (A) Schematic representation of the full-length Nnk protein. (B) AlphaFold3 prediction of the Nnk ZAD dimer. Protein structure was visualized using UCSF ChimeraX.^65^ pTM and iPTM values are used as proxies for the accuracy of predicting the overall protein folding and the relative positioning of the monomers within the dimer, respectively (hereinafter the same). (C) Western blot analysis of GFP-3xFLAG-tagged endogenous Nnk and exogenously expressed Nnk rescue constructs. Two-to-four hour embryos from corresponding homozygous strains were used for the analysis. As a control, 2-4 h *yw* embryos were used. (D) Airyscan imaging of GFP-3xFLAG-tagged WT and ΔZAD Nnk expressed under the control of the native *Nnk* promoter. Images were taken ∼15 min after entry into nc14. Embryos from corresponding homozygous strains were used for the analysis. The maximum intensity projected images are shown. (E) Schematic of the full-length CG30020 / Brother of Nicknack (BroN) protein. (F) AlphaFold3 prediction of the BroN ZAD dimer. Protein structure was visualized using UCSF ChimeraX.^65^ (G) Western blot analysis of GFP-3xFLAG-tagged endogenous BroN and exogenously introduced BroN rescue constructs. Two-to-four hour embryos from corresponding homozygous strains were used for the analysis. As a control, 2-4 h *yw* embryos were used. (H) Airyscan imaging of GFP-3xFLAG-tagged WT and ΔZAD BroN expressed under the control of the native *BroN* promoter. Images were taken ∼15 min after entry into nc14. Embryos from corresponding homozygous strains were used for the analysis. The maximum intensity projected images are shown. (I) Full-locus deletion alleles (*Nnk[Δ1]* and *BroN[Δ1]*) were produced by CRISPR/Cas9-mediated genome-editing. (J and K) ZAD-deficient transgenes failed to restore the lethal phenotype of homozygous *Nnk[Δ1]* (J) and *BroN[Δ1]* (K) mutants. See Figure S3.

To further characterize the *in vivo* functionalities of *Nnk* and *BroN* genes, we employed CRISPR/Cas9-mediated genome-editing to remove the entire transcription unit from the endogenous loci (Figure 2I: *Nnk[Δ1]* and *BroN[Δ1]* denote the resulting full-locus deletion alleles). Consistent with previous RNAi-mediated knockdown experiments,^7^ the homozygous *Nnk[Δ1]* mutant exhibited a complete lethal phenotype despite of its young evolutionary age (Figure 2J). Importantly, transgene rescue assays demonstrated that only the ZAD-containing WT construct was able to restore viability (Figure 2J), implicating that ZAD-mediated clustering activity is required for the *in vivo* function of Nnk. As in the case of *Nnk[Δ1]*, the homozygous *BroN[Δ1]* mutant also exhibited a complete lethal phenotype (Figure 2K), which was rescued only by the ZAD-containing WT construct (Figure 2K). Overall, these data are consistent with the idea that the correct functionalities of Nnk and BroN rely on the N-terminal ZAD, which has the ability to mediate condensate assembly in developing embryos.

### ZAD guides Nnk/BroN into H3K9me3-enriched heterochromatic regions

It has been previously reported that exogenously expressed Nnk and its paralogue Odj preferentially accumulate at peri-centromeric H3K9me3-enriched heterochromatic regions in S2 cultured cells,^7^ but their endogenous binding profiles in the context of *Drosophila* development are unknown. To address this point, we performed ChIP-seq analysis using a well-established anti-FLAG M2 antibody^47^ for all the genome-edited strains produced in this study (see Method Details), allowing us an unambiguous comparison of genome-wide distribution of ZAD-ZnFs and other insulator proteins by minimizing potential experimental noise arising from the variability of antibody quality for individual target proteins.^20,21^ Initially, to test the validity of our experimental approach, we first performed ChIP-seq analysis against endogenous CTCF and Su(Hw) using our newly produced genome-edited strains (Figure S2A-B). Comparison of our data with publicly available CTCF and Su(Hw) ChIP-seq profiles^43^ revealed essentially the same genome-wide distribution patterns of these proteins (Figure S2C-D), suggesting that GFP-3xFLAG tagging does not alter the binding profiles of these key DNA-binding proteins. We then performed ChIP-seq analysis of endogenous Nnk, BroN, and Odj in nc14 embryos by following the experimental procedure described in a previous study^43^ with minor modifications (see Method Details). Intriguingly, we found that the newly identified BroN globally colocalizes with Nnk and Odj (Figure 3A-D). These three proteins were found to be highly accumulated at H3K9me3-enriched heterochromatic regions defined by Cut&Tag analysis in nc14 embryos,^48^ but unexpectedly, the majority of peaks (∼70%) are present at H3K9me3-depleted regions (Figure 3A and C), suggesting that the H3K9me3 histone modification is not a prerequisite for the recruitment of these ZAD-ZnFs onto the genome. Analysis of ChIP-seq peak distribution revealed that Odj, Nnk, and BroN are mainly localized at transcription start sites (TSSs) and intragenic regions (Figure S3C). To elucidate the role of ZAD-mediated clustering activity *in vivo* (Figure 2D and H), the genome-wide distribution of ZAD-deficient Nnk and BroN mutants was further analyzed by ChIP-seq. Before proceeding with this experiment, we first confirmed that unmodified WT Nnk and BroN expressed as transgenes exhibit essentially the same ChIP-seq profiles as their endogenous counterparts in nc14 embryos (Figure S3D-E). Interestingly, peaks of Nnk and BroN mutants were found to be selectively lost from H3K9me3-enriched heterochromatic regions, while other peaks were largely maintained (Figure 3E-F, Figure S4A-C), suggesting that the N-terminal ZAD facilitates association of these proteins with H3K9me3-enriched heterochromatic regions in developing embryos. Given our preceding finding that ZAD-deficient mutants were unable to rescue the lethal phenotype of the full-locus deletion alleles produced in this study (Figure 2J-K), we suggest that ZAD-dependent recruitment of Nnk and BroN into H3K9me3-enriched heterochromatic regions is essential for the proper progression of *Drosophila* development. It is tempting to speculate that the N-terminal ZAD served as a key scaffold driving a co-evolutionary arms race between rapid alternations of heterochromatic regions and heterochromatin-interacting ZAD-ZnFs during insect evolution. ^7^

**Figure 3.**
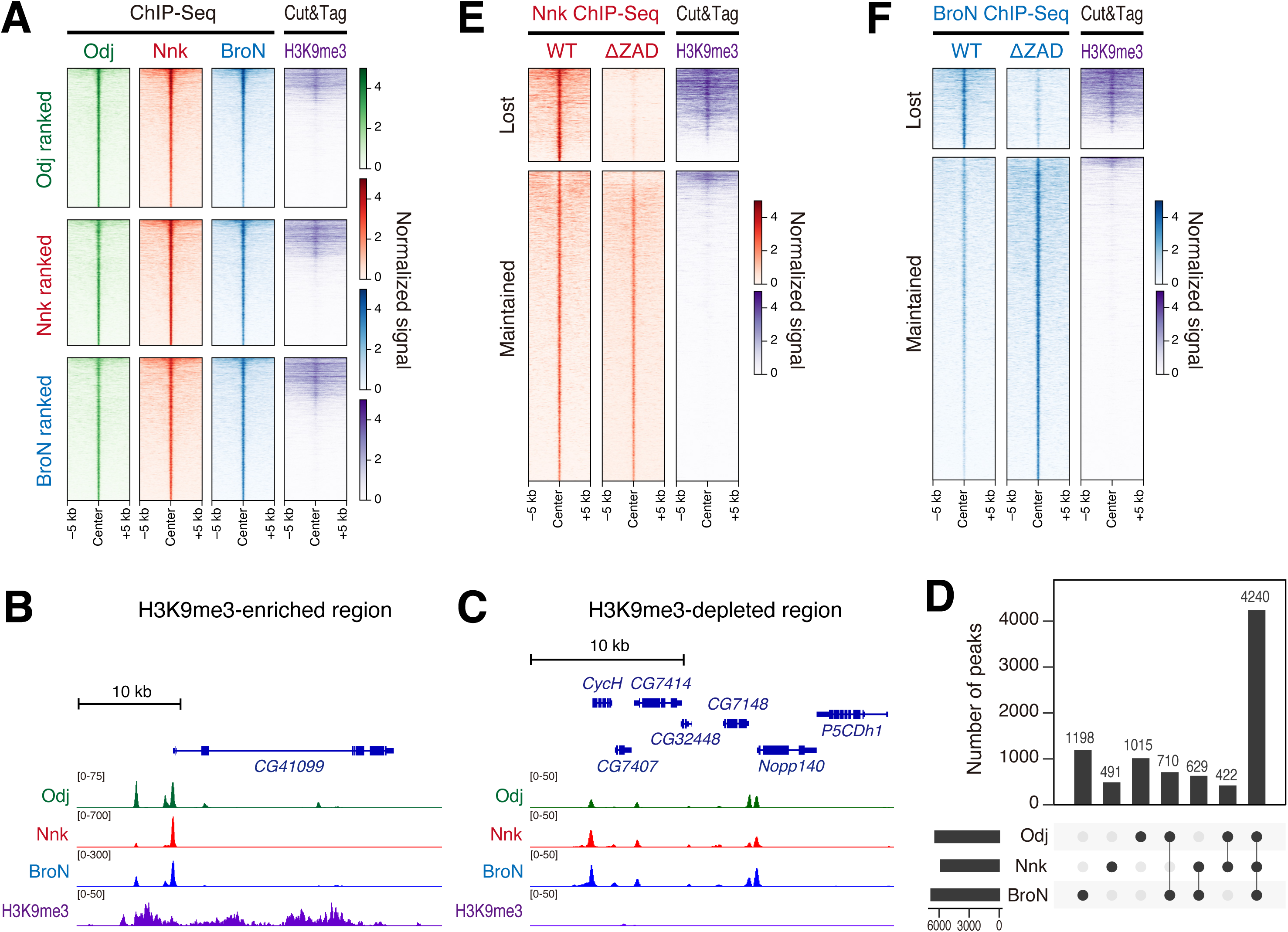
ZAD guides Nnk and BroN into H3K9me3-enriched repressive regions. (A) Heatmap visualization of Odj, Nnk, and BroN ChIP-seq peaks along with H3K9me3 Cut&Tag signal^48^ sorted by the strength of Odj (upper panels), Nnk (middle panel) or BroN (lower panels) signal. (B) IGV genome browser tracks showing Odj, Nnk, and BroN ChIP-seq coverage at the H3K9me3-enriched region. (C) IGV genome browser tracks showing Odj, Nnk, and BroN ChIP-seq coverage at the H3K9me3-depleted region. (D) UpSet plot showing the co-binding profiles of Odj, Nnk, and BroN. (E) Heatmap visualization of WT and ΔZAD Nnk along with H3K9me3 Cut&Tag signal^48^ sorted by the strength of WT Nnk. (F) Heatmap visualization of WT and ΔZAD BroN along with H3K9me3 Cut&Tag signal^48^ sorted by the strength of WT BroN. See Figure S3 and S4.

### ZAD-independent function of D19B during early embryogenesis

We next focused on the analysis of two neighboring paralogous genes, *D19A* and *D19B* (Figure 4A), as an example of non-clustering ZAD-ZnFs in developing embryos (Figure 1B). Western blot analysis revealed that D19B is more abundantly expressed than its paralogue D19A in early embryos (Figure 4B). To experimentally test their *in vivo* functions, CRISPR/Cas9-mediated genome-editing was employed to remove the entire transcription unit of *D19A* and *D19B* individually from the endogenous tandem locus (Figure 2I) (*D19A[Δ1]* and *D19B[Δ1]* denote the resulting full-locus deletion alleles). As a control, a recently reported ZAD-ZnF *CG2678/Kipf* locus^14^ was similarly engineered to remove the entire transcription unit (referred to as *kipf[Δ3]* since *kipf[Δ1]* and *kipf[Δ2]* alleles have already been described by Baumgartner and colleagues^14^). Contrary to *Nnk[Δ1]* and *BroN[Δ1]*, the resulting *D19A[Δ1]*, *D19B[Δ1]*, and *kipf[Δ3]* mutants were homozygous viable. We then measured the hatching rates of embryos collected from homozygous mutant adults. Consistent with a previous genetic study,^14^ the null mutation of *kipf* resulted in a reduction of embryo hatching rate (Figure 4C), presumably due to partial derepression of transposable elements. Similarly, there was a clear reduction in the hatching rate of embryos obtained from homozygous *D19A[Δ1]* mutant adults (Figure 4C). Strikingly, there was more than 90% reduction in the hatching rate when embryos obtained from homozygous *D19B[Δ1]* mutant adults were analyzed (Figure 4C), indicating that D19B plays an essential role in the correct progression of embryogenesis. Supporting this view, live visualization of His2Av-emiRFP670 in embryos obtained from homozygous *D19B[Δ1]* mutant adults revealed catastrophic nuclear defects, including asynchronous mitosis, disordered nuclei, and M phase arrest (Figure 4D). These severe negative effects were not seen when control *yw* virgin females were crossed with homozygous *D19B[Δ1]* mutant males (Figure 4E), indicating that the loss of maternal supply of D19B is responsible for the phenotype we observed. Given a recent finding that Kipf helps to protect genome integrity through the suppression of transposable elements by facilitating piRNA production in developing ovaries,^14^ we examined the possibility that D19A and D19B function in the same pathway. Consistent with the previous study, Kipf-GFP-3xFLAG formed bright foci within nurse cell nuclei that are thought to represent a subset of dual-strand piRNA clusters (Figure S5A-B). In contrast, both D19A and D19B showed very uniform nuclear distribution in nurse cell and surrounding follicle cell nuclei (Figure S5A-B). Importantly, derepression of Kipf-target transposable elements, *Burdock* and *3S18* (*bel*), was seen only in homozygous *kipf[Δ3]* ovaries in qRT-PCR analysis (Figure S5C). This result was further supported by RNA-seq analysis of dissected ovaries from homozygous *D19B[Δ1]* mutant females (Figure S5D-E). We therefore concluded that D19A and D19B act differently from Piwi-piRNA pathway-related ZAD-ZnF Kipf.^14^

**Figure 4.**
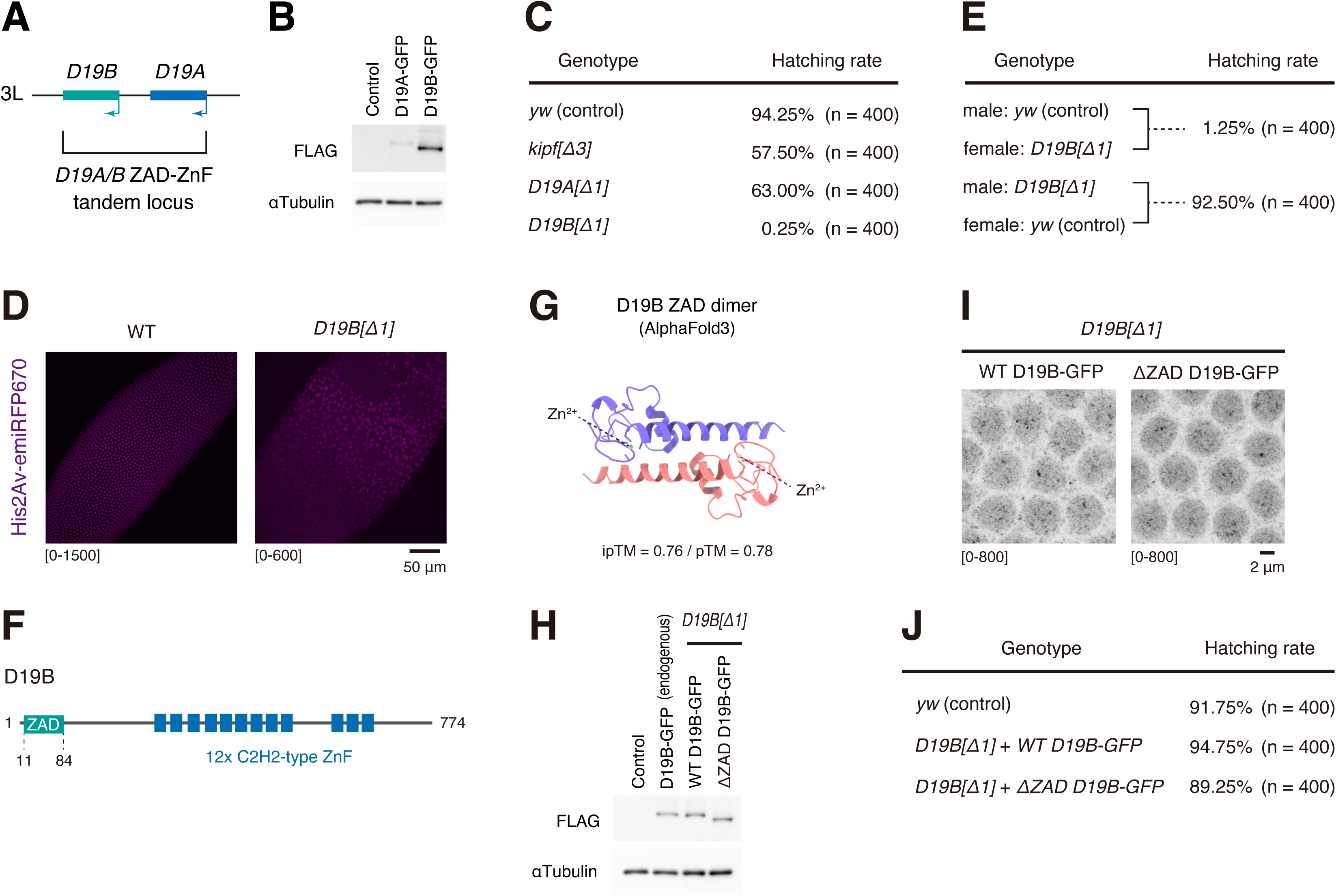
ZAD-independent function of D19B. (A) Two paralogous ZAD-ZnF genes *D19B* and *D19A* are tandemly placed at the locus. (B) Western blot analysis of GFP-3xFLAG-tagged endogenous D19A and D19B proteins. Two-to-four hour embryos from corresponding homozygous strains were used for the analysis. As a control, 2-4 h *yw* embryos were used. (C) Measurement of the embryo hatching rate. Embryos from corresponding homozygous full-locus deletion mutant adults were used for the analysis. (D) Confocal images of His2Av-emiRFP670 in embryos from WT or homozygous *D19B[Δ1]* mutant adults. The maximum intensity projected images of His2Av-emiRFP670 are shown. (E) Measurement of the embryo hatching rate. Maternal supply of D19B is essential for the correct progression of embryogenesis. (F) Schematic of the full-length D19B protein. (G) AlphaFold3 prediction of the D19B ZAD dimer. Protein structure was visualized using UCSF ChimeraX.^65^ (H) Western blot analysis of GFP-3xFLAG-tagged endogenous D19B and WT/ΔZAD D19B rescue constructs reintroduced into the endogenous locus via RMCE. Two-to-four hour embryos from corresponding homozygous mutant adults were used for the analysis. As a control, 2-4 h *yw* embryos were used. (I) Airyscan imaging of GFP-3xFLAG-tagged WT and ΔZAD D19B expressed from the endogenous locus. Images were taken ∼15 min after entry into nc14. Embryos from corresponding homozygous strains were used for the analysis. The maximum intensity projected images are shown. (J) Measurement of embryo hatching rate. Embryos from corresponding homozygous RMCE strains were used for the analysis. See Figure S4 and S5.

Motivated by our finding that Nnk and BroN exert their functions through the N-terminal ZAD domain (Figures 2 and 3), we then sought to elucidate the role of the D19B ZAD in detail (Figure 4F). As in the case of Nnk and BroN, AlphaFold3 predicted that the D19B ZAD also forms a dimer through its hydrophobic surface (Figure 4G). By using a recombinase-mediated cassette exchange (RMCE) approach,^49^ WT and ΔZAD rescue constructs were individually reintroduced into the endogenous *D19B[Δ1]* mutant allele. Western blot analysis showed that both rescue constructs are expressed at levels comparable to endogenous D19B (Figure 4H). Intriguingly, unlike Nnk and BroN (Figures 2D and H), both WT and ΔZAD D19B exhibited a very similar pattern of nuclear localization in developing embryos (Figure 4I). Moreover, measurement of embryo hatching rates revealed that the developmental defects seen in D19B-null embryos were almost fully restored by introducing either WT or ΔZAD D19B constructs (Figure 4J), indicating that D19B does not rely on the N-terminal ZAD to exert its function during embryogenesis. Overall, our data are consistent with the idea that each ZAD-ZnF has differential dependencies on the N-terminal ZAD in *Drosophila*.

### Comparative ChIP-seq analysis of two paralogous D19A and D19B proteins

To determine the distribution patterns of the two paralogous proteins D19A and D19B genome-wide, we next performed ChIP-seq analysis using the same standard M2 anti-FLAG antibody for both (see Method Details). As shown in the schematics (Figure 5A), D19A and its paralogue D19B share a very similar domain organization, including the number of zinc-finger domains and their amino acid composition (Figure 4F, Figure 5B), as well as the N-terminal ZAD dimerization scaffold (Figure 5C). Intriguingly, despite these extensive molecular similarities, we noticed that large fractions of D19A and D19B are recruited to distinct genomic locations without overlapping with H3K9me3, Nnk, and BroN peaks (Figure 5D-F, Figure S4D-E). These results imply that the two paralogous proteins D19A and D19B have functionally diverged after gene duplication during *Drosophila* evolution, with D19B playing a more dominant role than D19A in the early embryo (Figure 4C). To characterize the molecular function of the N-terminal D19B ZAD, we then performed ChIP-seq analysis of ΔZAD D19B expressed in the *D19B[Δ1]* mutant background. Consistent with our preceding hatching rate analysis (Figure 4J), D19B binding profiles were found to be mostly unchanged even in the absence of the N-terminal ZAD (Figure 5G-H, Figure S4F). Thus, we concluded that condensation-free D19B does not require the N-terminal ZAD dimerization surface to exert its function during embryogenesis, which sharply contrasts with the strict ZAD dependency of the condensate-forming Nnk and BroN (Figures 2 and 3).

**Figure 5.**
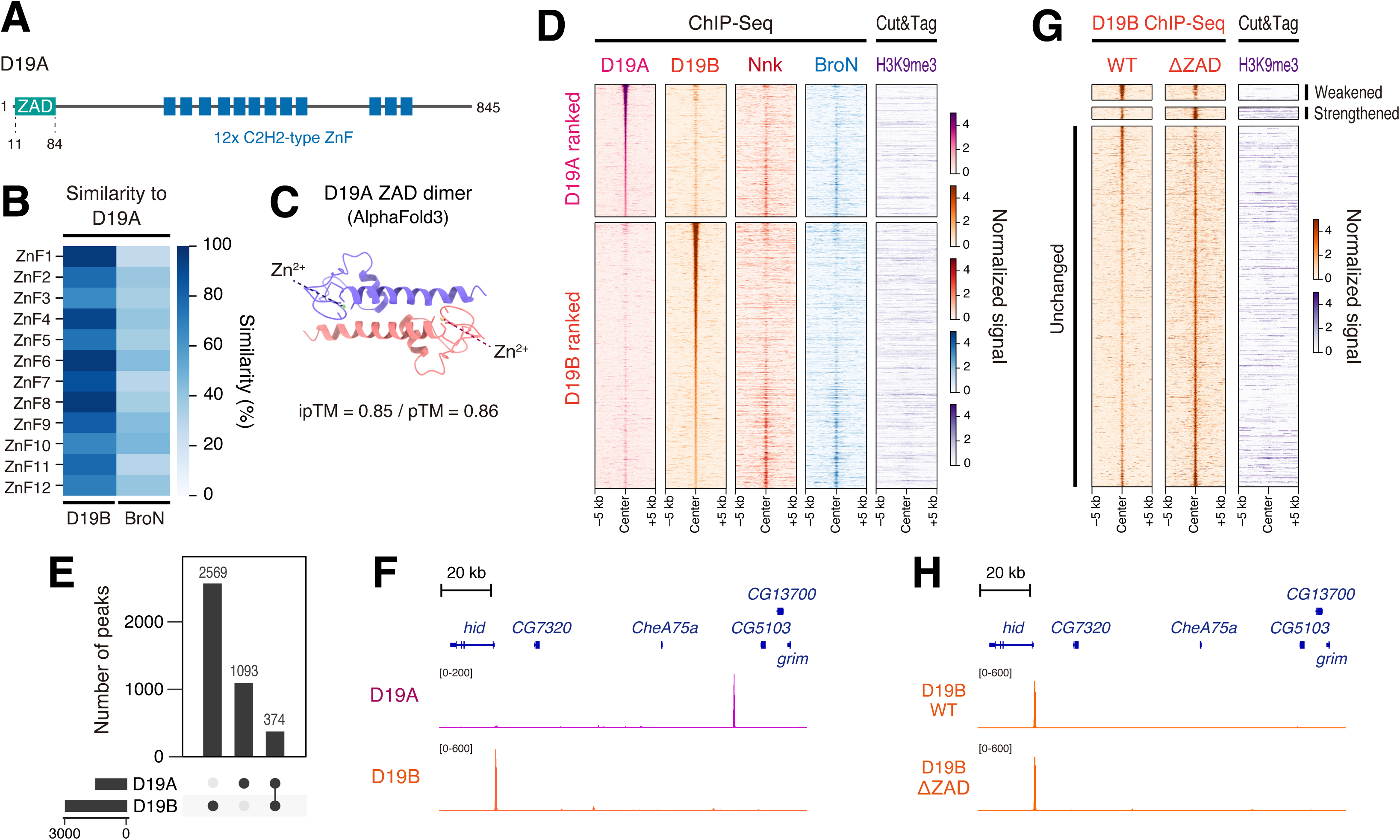
Functional divergence of two paralogous proteins D19A and D19B. (A) Schematic representation of the full-length D19A protein. (B) Similarity score calculated for individual zinc-finger domains of D19B and BroN relative to those of D19A. Zinc-finger domains of BroN were used as a negative control. (C) AlphaFold3 prediction of the D19A ZAD dimer. Protein structure was visualized using UCSF ChimeraX.^65^ (D) Heatmap visualization of D19A, D19B, Nnk, and BroN ChIP-seq distribution along with H3K9me3 Cut&Tag signal^48^ sorted by the strength of D19A (upper panels) or D19B (lower panels) ChIP-seq peaks. (E) UpSet plot showing the binding profiles of D19A and D19B. (F) IGV genome browser tracks showing D19A and D19B ChIP-seq coverage. (G) Heatmap visualization of WT and ΔZAD D19B ChIP-seq peaks. (H) IGV genome browser tracks showing WT and ΔZAD D19B ChIP-seq coverage. See Figure S5.

### Insulator binding activity is a common feature of ZAD-ZnFs

To systematically characterize the genome-wide distribution of ZAD-ZnFs in developing embryos, we performed ChIP-seq analysis for all the GFP-3xFLAG-tagged endogenous ZAD-ZnFs along with the non-ZAD-ZnF insulator proteins produced in this study (Figure 1, Figure S2). Intriguingly, integrative analysis of the obtained ChIP-seq profiles and publicly available Micro-C data of nc14 embryos^50^ revealed a preferential enrichment of ZAD-ZnFs at topological boundaries (Figure 6 and Figure S6A). For example, the *Abd-B*/*abd-A* Hox locus at the BX-C consists of a series of topologically associating domains (TADs) bordered by well-characterized insulator elements (*Fab-8*, *Fab-7*, *Fab-6*, *Mcp*, *Fab-4*, *Fab-3*, *Fub*) (reviewed in Kyrchanova *et al*.^51^). It was found that many of the ZAD-ZnFs, together with well-characterized core insulator proteins (*e.g.*, CP190, CTCF, BEAF-32), co-occupy these insulator elements (Figure 6A). Among these, enrichment of ZAD-ZnFs was more evident at sharp topological boundaries such as *Fub*, *Fab-6*, *Fab-8*, and the 5’ *Abd-B* TAD border (Figure 6A).^52^ Similar results were seen for the well-characterized insulator elements responsible for shaping 3D genome topology at the *ftz* and *eve* pair-rule loci (Figure 6B, Figure S6A).^53–57^ These data together suggest that ZAD-ZnFs have intrinsic insulator-binding activity in the early *Drosophila* embryo. Importantly, it also came to our attention that the tethering element that mediates long-range focal contact at the *ftz*/*Scr* locus^50^ (Figure 6B; arrowhead) is also enriched with many ZAD-ZnFs (Figure 6B; orange region). It is tempting to speculate that not only GAF^58^ but also ZAD-ZnFs play a role in organizing long-range focal interactions in developing embryos (see Discussion).

**Figure 6.**
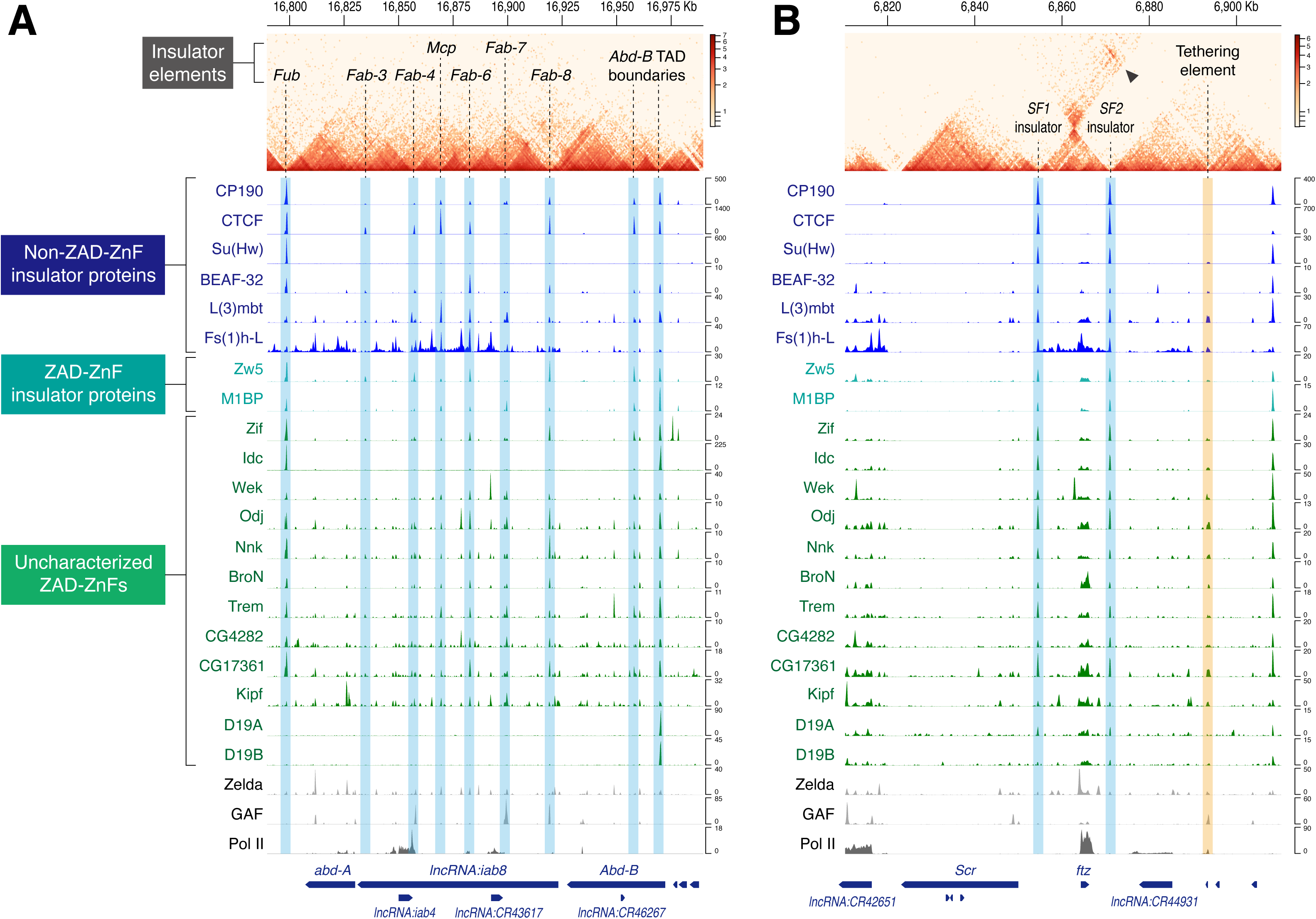
Topological boundaries are associated with ZAD-ZnF cluster. (A) Coolbox toolkit^66^ was used to visualize Micro-C map along with ChIP-seq profiles of ZAD-ZnFs and non-ZAD-ZnF insulator proteins at the *Abd-B/abd-A* Hox locus. Publicly available Micro-C data in nc14 embryos (Batut *et al.*^50^ ; SRR14136009) was used for the analysis. (B) Coolbox toolkit^66^ was used to visualize Micro-C map along with ChIP-seq profiles of ZAD-ZnFs and non-ZAD-ZnF insulator proteins at the *Scr/ftz* Hox locus. Publicly available Micro-C data in nc14 embryos (Batut *et al.*^50^ ; SRR14136009) was used for the analysis. See Figure S6.

### ZAD-ZnFs colocalize with known insulator proteins genome-wide

To further address the degree of co-occupancy between ZAD-ZnFs and non-ZAD-ZnF core insulator proteins genome-wide, we analyzed the distribution profiles of ZAD-ZnFs using the ChIP-seq peaks of BEAF-32, CTCF, Su(Hw), and CP190 as viewpoints. Consistent with a previous study in nc14 embryos,^43^ the differential distribution patterns of BEAF-32, CTCF, and Su(Hw) were nicely reproduced in our hands (Figure 7A), verifying the quality of our ChIP-seq dataset. Having confirmed this, we next visualized the distribution of all tested ZAD-ZnFs at the four viewpoints. Intriguingly, it turned out that ZAD-ZnFs globally co-localize with BEAF-32, CP190, and CTCF in developing *Drosophila* embryos (Figure 7B-C, Figure 6B). Co-localization of ZAD-ZnFs and Su(Hw) was also seen genome-wide, but the strong Su(Hw) peaks tended to be depleted of binding of ZAD-ZnFs (Figure S6C), as well as other insulator proteins (Figure 7A). Importantly, the two paralogous ZAD-ZnFs D19A and D19B did not show clear enrichment at the peaks of any of the four viewpoint insulator proteins genome-wide (Figure 7B-C, Figure S6B-C), implicating that they are recruited to only a specific subset of topological boundaries, such as the 5’ *Abd-B* TAD border and the *SF2* insulator (Figure 6), to exert their locus-specific functions. Overall, such an extensive degree of genome-wide colocalization with core insulator proteins suggests that the diverse molecular functions of ZAD-ZnFs have evolutionarily arisen from their ancestral role as insulator-binding proteins.

**Figure 7.**
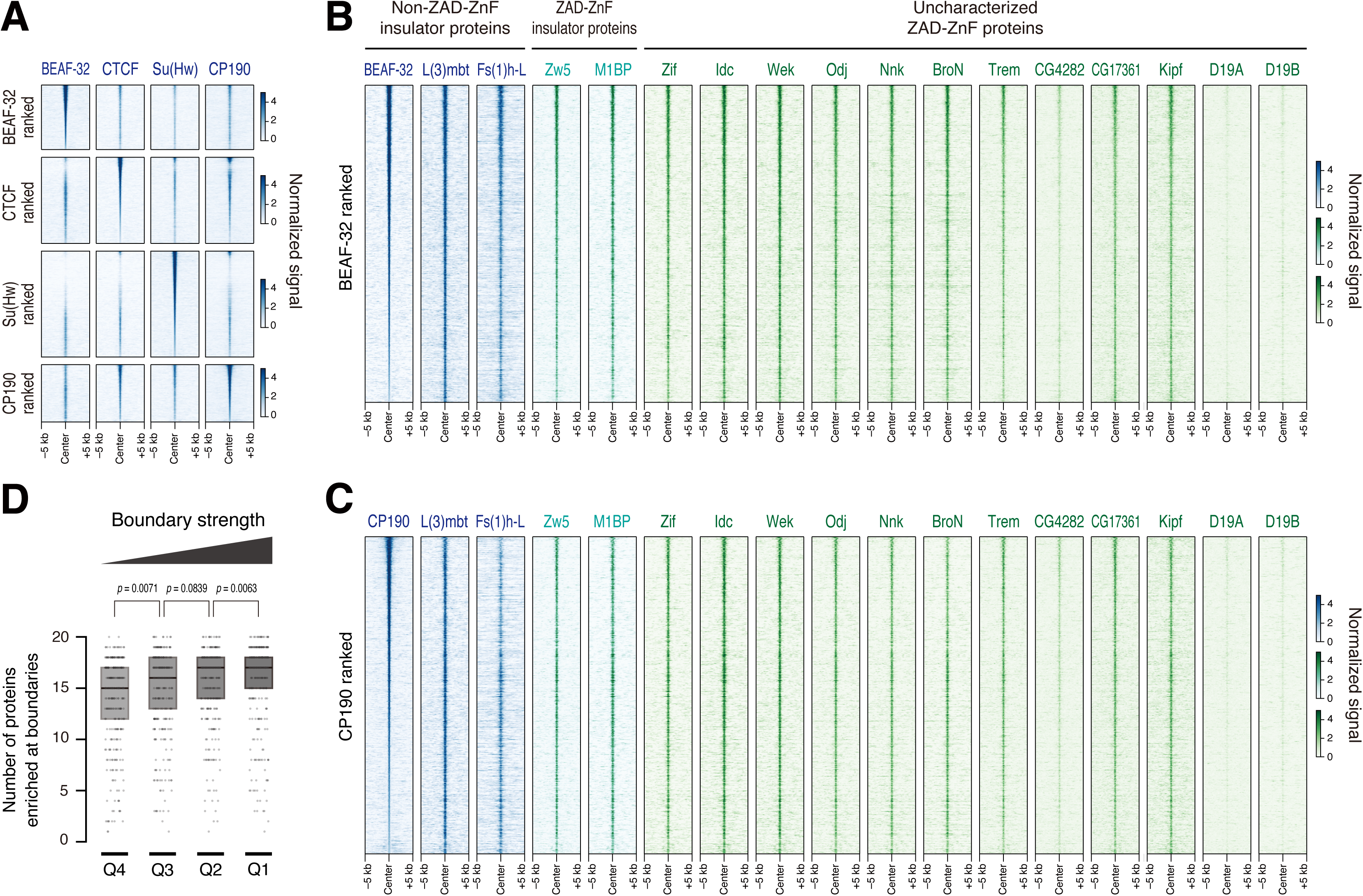
Intrinsic insulator binding activity of ZAD-ZnFs. (A) Heatmap visualization of BEAF-32, CTCF, Su(Hw), and CP190 ChIP-seq distribution sorted by the strength of BEAF-32 (upper panels), CTCF (upper middle panel), Su(Hw) (lower middle panel) or CP190 (lower panels) ChIP-seq peaks. (B) Heatmap visualization of ChIP-seq distribution of non-ZAD-ZnF insulator proteins and ZAD-ZnFs sorted by the strength of BEAF-32 ChIP-seq peaks. (C) Heatmap visualization of ChIP-seq distribution of non-ZAD-ZnF insulator proteins and ZAD-ZnFs sorted by the strength of CP190 ChIP-seq peaks. (D) Topological boundaries are divided into quantiles according to the boundary strength^50^. The numbers of proteins containing ChIP-seq peaks were determined for individual boundaries. List of proteins used for the analysis is shown in Figure S7B. In total, 336 (Q1), 336 (Q2), 337 (Q3), and 337 (Q4) boundaries were analyzed. p-values of two-sided Wilcoxon rank-sum test are shown. See Figure S6 and S7.

Next, to address the mechanism underlying genome-wide co-occupancy of ZAD-ZnFs with other insulator proteins, we analyzed the ChIP-seq profiles of the ZAD-deficient Nnk and BroN mutants (Figure 2). It turned out that both Nnk and BroN retain co-localization with BEAF-32 and CP190 genome-wide, even in the absence of the N-terminal ZAD and the accompanying condensation activity (Figure 2D and H, Figure S7A), suggesting that the C-terminal zinc-finger arrays, but not the N-terminal ZAD, are responsible for their association with insulator elements. Lastly, to explore the functional link between boundary strength and the ChIP-seq binding profiles of ZAD-ZnFs, we calculated the number of proteins containing peaks at individual topological boundaries (Figure S7B). This analysis revealed that there is a positive correlation between boundary strength and the number of proteins associating with the corresponding DNA regions (Figure 7D), implicating that the co-occupancy of ZAD-ZnFs and other non-ZAD-ZnF insulator proteins fosters the establishment and/or maintenance of topological boundaries during early embryogenesis (see Discussion).

## Discussion

### ZAD-ZnFs as insulator binding proteins

A striking finding of this study is that not only the previously characterized Zw5 and M1BP^16,17^ but also other ZAD-ZnFs are highly accumulated at the topological boundaries of key developmental loci in developing *Drosophila* embryos (Figure 6, Figure S6A). Analysis of ChIP-seq datasets further revealed extensive co-occupancy of ZAD-ZnFs and other non-ZAD-ZnF insulator proteins genome-wide (Figure 7, Figure S6B-C), giving rise to the possibility that clustered binding of ZAD-ZnFs helps to organize 3D genome topology. Given recent findings that the majority of topological boundaries are maintained even after the depletion of well-characterized core insulator proteins such as CTCF, BEAF-32, and CP190,^43,59^ it is tempting to speculate that insulator-associating ZAD-ZnFs help to ensure the robustness of topological boundary formation in concert with other insulator proteins during *Drosophila* development. In mammals, ubiquitously expressed DNA-binding proteins YY1 and Ldb1 are suggested to mediate long-range interactions through homotypic protein-protein interactions mediated by their dimerization domains.^60,61^ Similarly in *Drosophila*, it was recently shown that the dimerization activity of the N-terminal POZ/BTB domain of GAF can mediate long-range interactions over large genomic distances.^58^ In analogy to the actions of YY1, Ldb1, and GAF, it can be possible that ZAD-ZnFs use their dimerization activity to help establish topological boundaries through homotypic protein-protein interactions. This mechanism might also be at work during the establishment of long-range focal interactions mediated by tethering elements (Figure 6B; orange region).^50^ Importantly, our data suggest that the N-terminal ZAD is dispensable for the recruitment of ZAD-ZnFs to insulator elements (Figure S7A), but this does not necessarily rule out the possibility that the dimerization activity of the N-terminal ZAD helps organize higher-order genome topology after loading onto insulator elements. Clearly, future studies are needed to fully elucidate the molecular functions of individual insulator-associating ZAD-ZnFs during the formation of 3D genome topology in *Drosophila* and other insect species.

### Heterochromatin association of ZAD-ZnFs

It has been previously reported that exogenously expressed Nnk and Odj are highly accumulated at H3K9me3-enriched heterochromatic regions in S2 cultured cells.^7^ Consistent with this observation, our ChIP-seq analysis of endogenous Nnk and Odj revealed their preferential association with H3K9me3-enriched heterochromatic regions in developing embryos (Figure 3). We further demonstrated that a newly identified CG30020/BroN cooperates with Nnk and Odj to associate with their target sites across the genome (Figure 3). While the detailed molecular functions of these H3K9me3-associating ZAD-ZnFs remain to be elucidated at this point, our data indicate that ZAD-dependent condensation activity is required for their correct *in vivo* functionalities (Figure 2). Intriguingly, it has been previously suggested that phase separation of HP1α mediates dynamic compaction of heterochromatic regions to silence gene expression in both mammals and *Drosophila*.^62,63^ Similarly, *Drosophila* GAF subnuclear foci have been reported to contribute to the silencing of heterochromatic AAGAG satellite repeats^28^. Given these findings, it is conceivable that ZAD-dependent co-condensation of Nnk, Odj, and BroN helps to ensure gene silencing at heterochromatic regions in developing embryos. This mechanism might involve the physical isolation of heterochromatic regions into repressive ZAD-ZnF condensates, thereby excluding active transcriptional machinery from silenced loci. A similar exclusion mechanism has been suggested to operate in PRC1-mediated gene silencing in mammals.^64^ Strict ZAD-dependency of H3K9me3-association activities of Nnk and BroN (Figure 3) gives rise to the possibility that the N-terminal ZAD served as a key scaffold driving a co-evolutionary arms race between rapid alternations of heterochromatic regions and heterochromatin-interacting ZAD-ZnFs during insect evolution. ^7^

### Dynamic modulation of ZAD-ZnF functions

While all the ZAD-ZnFs share a common structural arrangement (*i.e.*, an N-terminal ZAD followed by an array of zinc-finger domains at the C-terminus), only a subset of ZAD-ZnFs exhibits condensation activity in living embryos (Figure 1B). This implicates that the dimerization activity of the N-terminal ZAD is not sufficient to mediate the local accumulation of ZAD-ZnFs within a nucleus. Importantly, Kipf forms very bright foci that are thought to represent a subset of dual-strand piRNA clusters in nurse cell nuclei (Figure S5A-B), but its nuclear distribution is rather uniform in early embryos (Figure 1B), suggesting that the protein localization of ZAD-ZnFs can be dynamically modulated during the progression of *Drosophila* development. Given a recent finding that the interaction between Kipf and Rhino, a germline-specific variant of HP1, is responsible for the formation of bright Kipf foci in nurse cell nuclei,^14^ we speculate that the presence or absence of binding partners of individual ZAD-ZnFs can flexibly tune their genome-wide distribution and *in vivo* functionalities. It is conceivable that the gain or loss of binding partners has also contributed to the molecular diversification and functional specification of evolutionarily dynamic ZAD-ZnFs in other insect species. As a future study, it would be intriguing to explore the roles of individual ZAD-ZnFs in developmental contexts other than early embryos. We believe that our current study represents a critical starting point toward a comprehensive understanding of the diverse, multifaceted functionalities of ZAD-ZnFs in *Drosophila* and other insect species.

## Acknowledgement

We thank Hitomi Takishita and Misako Sato for fly husbandry, and the Bloomington *Drosophila* Stock Center for fly strains. We are also grateful to Yusuke Kishi, Yuka W. Iwasaki, Chikara Takeuchi and Ryuichiro Nakato for their advice on ChIP-seq experiment and members of the Fukaya laboratory for critical comments on the manuscript. T.F. was supported by JST FOREST program (JPMJFR214W), the Grant-in-Aid for Scientific Research (A) (25H00967), the Grant-in-Aid for Transformative Research Areas (A) (24H02327) from JSPS, and the research grant from the Takeda Science Foundation. R.S. was supported by AMED ASPIRE (JP23jf0126003) and the Grant-in-Aid for Young Scientists (25K18438) from JSPS. Y.U. was supported by JSPS fellowship (24KJ0843). S.M. was supported by a Grant-in-Aid for Scientific Research (A) (23K05631) from the JSPS and the research grants from the Sumitomo Foundation, Nakajima Foundation, and Inamori Foundation.

## Author contributions

R.S. performed all the ChIP-seq analysis. Y.U. and T.F. performed the hatching rate analysis. S.M. and Y.U. performed the RNA-seq analysis. Y.U. performed the Micro-C data analysis. T.F. performed the CRISPR/Cas9-mediated genome-editing, the fly genetics, and the imaging analysis. T.F. supervised the project and wrote the manuscript. All the authors edited and approved the manuscript.

## Key Resources Table

| REAGENT or RESOURCE | SOURCE | IDENTIFIER |
| --- | --- | --- |
| **Antibodies** |  |  |
| Monoclonal Anti-FLAG® M2 antibody (mouse) | Sigma-Aldrich | Cat# F3165<br>RRID: AB_259529 |
| Monoclonal Anti-FLAG® M2 antibody (mouse) | Sigma-Aldrich | Cat# F1804<br>RRID: AB_262044 |
| Monoclonal Anti-α-Tubulin antibody (mouse) | Sigma-Aldrich | Cat# T5168<br>RRID: AB_477579 |
| Peroxidase AffiniPure Rabbit Anti-Mouse IgG (H+L) | Jackson ImmunoResearch | Cat# 315-035-003<br>RRID: AB_2340061 |
| Anti-RNA polymerase II monoclonal antibody CTD4H8 (mouse) | Merck Millipore | Cat# 05-623<br>RRID: AB_10013820 |
| **Bacterial and virus strains** |  |  |
| DH5α Competent Cells | This study | N/A |
| **Biological samples** |  |  |
| **Chemicals, peptides, and recombinant proteins** |  |  |
| Modified Silicone Fluids | Shin-Etsu Silicone | FL-100-450CS |
| Modified Silicone Fluids | Shin-Etsu Silicone | FL-100-1000CS |
| polylap (polyethylene membrane) | Ube Film | N/A |
| **Critical commercial assays** |  |  |
| TRIzol | Thermo Fisher | Cat# 15596026 |
| TB Green Premix Ex Taq II | Takara | Cat# RR820 |
| PrimeScript RT reagent kit with gDNA eraser | Takara | Cat# RR047 |
| EDTA-free protease inhibitor | Nacalai Tesque | Cat# 03969-34 |
| Dynabeads Protein G for Immunoprecipitation | Invitrogen | Cat# 10009D |
| NEBNext Ultra II DNA Library Prep kit for Illumina | New England Biolabs | Cat# E7645S |
| NEBNext Multiplex Oligos for Illumina (Index Primers Set 1 | New England Biolabs | Cat# E7335L |
| NEBNext Multiplex Oligos for Illumina (Index Primers Set 2 | New England Biolabs | Cat# E7500L |
| NEBNext Multiplex Oligos for Illumina (Index Primers Set 3 | New England Biolabs | Cat# E7710L |
| NEBNext Multiplex Oligos for Illumina (Index Primers Set 4 | New England Biolabs | Cat# E7730L |
| AMPure XP Reagent | Beckman Coulter | Cat# A63881 |
| TruSeq Stranded mRNA Library Prep | illumina | Cat# 20020594 |
| RNAClean XP | Beckman Coulter | Cat# A63987 |
| Deposited data |  |  |
| Raw and processed data (ChIP-seq) | This study | GEO: GSE295515 |
| Raw and processed data (RNA-seq) | This study | GEO: GSE295515 |
| CTCF ChIP-seq | Cavalheiro et al.<sup>43</sup> | http://furlonglab.embl.de/data |
| Su(Hw) ChIP-seq | Cavalheiro et al.<sup>43</sup> | http://furlonglab.embl.de/data |
| RNA-seq for *kipferl*-depleted and control ovaries | Baumgartner et al.<sup>14</sup> | GEO: GSE202468 |
| Micro-C for *yw* embryos | Batut et al.<sup>50</sup> | GEO: GSE171396 |
| Experimental models: Cell lines |  |  |
| Experimental models: Organisms/strains |  |  |
| *D. melanogaster*. *y*<sup>1</sup> *w*<sup>67c23</sup> *P{Crey}1b*; *sna*<sup>Sco</sup>/*CyO* | Bloomington Drosophila Stock Center | BDSC # 766 |
| *D. melanogaster*. *y*<sup>1</sup> *w*<sup>67c23</sup> *P{Crey}1b*; *D\*/TM3*, *Sb*<sup>1</sup> | Bloomington Drosophila Stock Center | BDSC # 851 |
| *D. melanogaster*. *y*[1] *M{vas-int.Dm}ZH-2A w[\*]*; *PBac{y[+]-attP-3B}VK00033* | Bloomington Drosophila Stock Center | BDSC # 24871 |
| *D. melanogaster*. *PBac{y[+]-attP-3B}VK00002* | Bloomington Drosophila Stock Center | BDSC # 9723 |
| *D. melanogaster*. *zelda-GFP-3xFLAG* | Hamamoto et al.<sup>67</sup> | N/A |
| *D. melanogaster*. *CTCF-GFP-3xFLAG* | This study | N/A |
| *D. melanogaster. Cp190-GFP-3xFLAG* | This study | N/A |
| *D. melanogaster. BEAF-32-GFP-3xFLAG* | This study | N/A |
| *D. melanogaster. fs(1)h-L-GFP-3xFLAG* | This study | N/A |
| *D. melanogaster. vasa-mCherry-3xFLAG* | This study | N/A |
| *D. melanogaster. Trl-GFP-3xFLAG* | This study | N/A |
| *D. melanogaster. Trl-mCherry-3xFLAG* | This study | N/A |
| *D. melanogaster. l(3)mbt-GFP-3xFLAG* | This study | N/A |
| *D. melanogaster. su(Hw)-GFP-3xFLAG* | This study | N/A |
| *D. melanogaster. Odj-GFP-3xFLAG* | This study | N/A |
| *D. melanogaster. Nnk-GFP-3xFLAG* | This study | N/A |
| *D. melanogaster. CG4282-GFP-3xFLAG* | This study | N/A |
| *D. melanogaster. dwg-GFP-3xFLAG* | This study | N/A |
| *D. melanogaster. D19A-GFP-3xFLAG* | This study | N/A |
| *D. melanogaster. D19B-GFP-3xFLAG* | This study | N/A |
| *D. melanogaster. M1BP-GFP-3xFLAG* | This study | N/A |
| *D. melanogaster. Zif-GFP-3xFLAG* | This study | N/A |
| *D. melanogaster. wek-GFP-3xFLAG* | This study | N/A |
| *D. melanogaster. trem-GFP-3xFLAG* | This study | N/A |
| *D. melanogaster. kipf-GFP-3xFLAG* | This study | N/A |
| *D. melanogaster. idc-GFP-3xFLAG* | This study | N/A |
| *D. melanogaster. BroN-GFP-3xFLAG* | This study | N/A |
| *D. melanogaster. CG17361-GFP-3xFLAG* | This study | N/A |
| *D. melanogaster. D19A[Δ1]* | This study | N/A |
| *D. melanogaster. D19B[Δ1]/tm3, Kr-GFP[w<sup>+</sup>]* | This study | N/A |
| *D. melanogaster. kipf[Δ3]* | This study | N/A |
| *D. melanogaster. Nnk[Δ1]/tm3, Kr-GFP[w<sup>+</sup>]* | This study | N/A |
| *D. melanogaster. BroN[Δ1]/CyO, Kr-GFP[w<sup>+</sup>]* | This study | N/A |
| *D. melanogaster. D19B RMCE[WT]* | This study | N/A |
| *D. melanogaster. D19B RMCE[ΔZAD]* | This study | N/A |
| *D. melanogaster. Nnk<sup>WT</sup>-GFP-3xFLAG (VK0002)* | This study | N/A |
| *D. melanogaster. Nnk<sup>ΔZAD</sup>-GFP-3xFLAG (VK0002)* | This study | N/A |
| *D. melanogaster. BroN<sup>WT</sup>-GFP-3xFLAG (VK0002)* | This study | N/A |
| *D. melanogaster. BroN<sup>ΔZAD</sup>-GFP-3xFLAG (VK0002)* | This study | N/A |
| *D. melanogaster. Nnk<sup>WT</sup>-GFP-3xFLAG; Nnk[Δ1]* | This study | N/A |
| *D. melanogaster. Nnk<sup>ΔZAD</sup>-GFP-3xFLAG; Nnk[Δ1]/tm3, Kr-GFP[w<sup>+</sup>]* | This study | N/A |
| *D. melanogaster. BroN<sup>WT</sup>-GFP-3xFLAG, BroN[Δ1]* | This study | N/A |
| *D. melanogaster. BroN<sup>ΔZAD</sup>-GFP-3xFLAG, BroN[Δ1]/CyO, Kr-GFP[w<sup>+</sup>]* | This study | N/A |
| *D. melanogaster. SV40NLS-GFP-3xFLAG (VK00033)* | This study | N/A |
| *D. melanogaster. His2Av-emiRFP670 (VK00028)* | This study | N/A |
| *D. melanogaster. His2Av-emiRFP670, D19B[Δ1]/tm3, Kr-GFP[w<sup>+</sup>]* | This study | N/A |
| Oligonucleotides |  |  |
| DNA oligos are listed in Table S1. | This study | N/A |
| Recombinant DNA |  |  |
| Plasmids are listed in Method Details. | This study | N/A |
| Software and algorithms |  |  |
| Fiji | Schindelin et al.<sup>68</sup> | www.imagej.nih.gov/ij |
| AlphaFold3 | Abramson et al.<sup>46</sup> | https://golgi.sandbox.google.com/about |
| UCSF ChimeraX | Pettersen et al.<sup>65</sup> | https://www.cgl.ucsf.edu/chimerax/ |
| fastp v0.23.4 | Chen et al.<sup>69</sup> | https://github.com/OpenGene/fastp |
| bowtie2 v2.4.1 | Langmead et al.<sup>70</sup> | https://github.com/BenLangmead/bowtie2 |
| samtools v1.13 | Li et al.<sup>71</sup> | https://github.com/samtools/samtools |
| macs2 v2.2.6 | Zhang et al.<sup>72</sup> | https://hbctraining.github.io/Intro-to-ChIPseq/lessons/05_peak_calling_mac.html |
| bedtools v2.30.0 | Quinlan and Hall.<sup>73</sup> | https://bedtools.readthedocs.io/en/latest/ |
| deeptools v3.5.1 | Ramírez et al.<sup>74</sup> | https://github.com/deeptools/deepTools |
| R v4.4.3 |  | https://www.R-project.org/ |
| Python v3.11.5 |  | https://www.python.org |
| GenomicRanges v1.22.4 | Lawrence et al.<sup>75</sup> | https://github.com/Bioconductor/GenomicRanges |
| UpSetR v1.4.0 | Conway et al.<sup>76</sup> | https://github.com/hms-dbmi/UpSetR |
| MAnorm v1.3.0 | Shao et al.<sup>77</sup> | https://github.com/shao-lab/MAnorm |
| Integrative Genome Viewer v2.18.0 | Robinson et al.<sup>78</sup> | https://igv.org |
| CoolBox v0.3.7 | Xu et al.<sup>66</sup> | https://github.com/GangCaoLab/CoolBox |
| DROMPA3 v3.7.2 | Nakato et al.<sup>79</sup> | https://github.com/rnakato/DROMPA3 |
| SSP v1.4.0 | Nakato et al.<sup>80</sup> | https://github.com/rnakato/SSP |
| Biopython v1.85 | Cock et al.<sup>81</sup> | https://biopython.org |
| STAR v2.7.10b | Dobin et al.<sup>82</sup> | https://github.com/alexdobin/STAR |
| TEcount v2.2.3 | Jin et al.<sup>83</sup> | https://github.com/mhammell-laboratory/TEtranscripts |
| DESeq2 v1.44.0 | Love et al.<sup>84</sup> | https://www.biocconductor.org/packages/release/bioc/vignettes/DESeq2/inst/doc/DESeq2.html |
| BWA v0.7.18 | Li et al.<sup>85</sup> | https://bio-bwa.sourceforge.net |
| pairtools v0.2.2 | Open2C, Abdennur et al.<sup>86</sup> | https://github.com/open2c/pairtools |
| Cooler v0.10.2 | Abdennur et al.<sup>87</sup> | https://github.com/open2c/cooler |
| Other |  |  |
| Zeiss LSM900 with Airyscan2 | Zeiss | N/A |
| Evident FLUOVIEW FV4000 | Evident | N/A |

## Resource Availability

### Data and code availability

All sequencing data have been deposited in the Gene Expression Omnibus under accession code GEO: GSE295515.

### Experimental Model and Subject Details

In all experiments, *Drosophila melanogaster* embryos or ovaries were analyzed.

## Method Details

### Site-specific transgenesis by phiC31 system

GFP-3xFLAG-fusion Nnk and BroN rescue constructs were integrated into a unique *attP* landing site on the second chromosome using VK00002 strain.^88^ SV40NLS-GFP-3xFLAG and His2Av-emiRFP670 transgenes were integrated into a unique *attP* landing site on the third chromosome using VK00033 strain and the second chromosome using VK00028, respectively.^88^ Microinjection was performed as previously described.^89^ In brief, 0-1 h embryos were collected and dechorionated with bleach. Aligned embryos were dried with silica gel for ∼8-9 min and covered with FL-100-1000CS silicone oil (Shin-Etsu Silicone). Subsequently, microinjection was performed using FemtoJet (Eppendorf) and DM IL LED inverted microscope (Leica) equipped with M-152 Micromanipulator (Narishige). Injection mixture typically contains ∼900 ng/µl plasmid DNA, 5 mM KCl, 0.1 mM phosphate buffer, pH 6.8. *mini-white* marker was used for screening.

### CRISPR/Cas9-mediated genome-editing

Genome-edited fly lines were obtained by CRISPR/Cas9-mediated homology directed repair. For GFP-3xFLAG/mCherry-3xFLAG insertion, gRNA was designed to target PAM-containing sequence adjacent to the stop codon. pCFD3 gRNA expression plasmid, pBS-Hsp70-Cas9 plasmid (addgene# 46294) and pBS-GFP-3xFLAG/mCherry-3xFLAG-dsRed-loxP donor plasmid containing 5’ and 3’ homology arms were co-injected using the standard *yw* laboratory strain. Injection mixture typically contains ∼300 ng/µl pCFD3 gRNA expression plasmid, ∼600 ng/µl pBS-Hsp70-Cas9 plasmid and ∼300 ng/µl donor plasmid. Microinjection was performed as described above. *3xP3-dsRed* was used for screening. Resulting genome-edited flies were then crossed with *y*^1^ *w*^67c23^ *P{Crey}1b; D*\**/TM3, Sb*^1^ (BDSC# 851) or *y*^1^ *w*^67c23^ *P{Crey}1b; sna^Sco^/CyO* (BDSC# 766) to remove *3xP3-dsRed* marker from the endogenous locus. For full locus deletion, two gRNAs were designed to target PAM-containing sequences upstream of the TSS and downstream of the stop codon. A pair of pCFD3 gRNA expression plasmids, pBS-Hsp70-Cas9 plasmid and pBS-3xFLAG-attP-dsRed-attP donor plasmid containing 5’ and 3’ homology arms were co-injected using the standard *yw* laboratory strain. Injection mixture typically contains ∼200 ng/µl pCFD3 gRNA expression plasmids, ∼400 ng/µl pBS-Hsp70-Cas9 plasmid and ∼300 ng/µl donor plasmid. *3xP3-dsRed* was used for screening. Deletion and insertion were confirmed by the PCR analysis of purified genomic DNA. DNA oligos used to build gRNA expression plasmids are listed in Table S1.

### Recombinase-mediated cassette exchange (RMCE)

D19B rescue constructs and p3xP3-EGFP.vas-int.NLS plasmid (addgene# 60948) were co-injected into embryos obtained from the *D19B[Δ1]/tm3, Kr-GFP[w+]* strain. Injection mixture typically contains ∼500 ng/µl rescue plasmid and ∼800 ng/µl phiC31 expression plasmid. Loss of *3xP3-dsRed* marker was used for screening.

### Western blotting

Twenty of 2-4 h embryos were hand collected and 50 μl of 2x SDS sample buffer was added. Subsequently, embryos were crushed with a pestle and boiled at 94 °C for ∼3 min. Insoluble pellet was removed by centrifugation and lysates were separated by SDS-PAGE, transferred to Immobilon-p PVDF membranes (Merck, IPVH0001), and immunoblotted with antibodies. The primary antibodies used were as follows: mouse monoclonal anti-FLAG M2 antibody (Sigma, F3165, 1:10000), mouse monoclonal anti-α-Tubulin antibody (Sigma, T5168, 1:40000). The membranes were then incubated with secondary antibodies conjugated with horseradish peroxidase (HRP). Chemiluminescence was detected by a FUSION SOLO.7S. EDGE (Vilber-Lourmat).

### Confocal imaging analysis

In Figure S1C and S5A-B, virgin female flies were collected and aged for 2 days before ovary dissection. Ovaries were dissected into 1x PBS buffer and mounted between a polyethylene membrane (Ube Film) and a coverslip (18 mm x 18 mm) and embedded in FL-100-450CS (Shin-Etsu Silicone). Dissected ovaries were imaged by a FV4000 confocal microscope equipped with SilVIR detector (Evident). UPlanXApo 40x / 1.4 N.A. oil immersion objective was used. Fluorescence of mCherry and GFP were excited using 561-nm and 488-nm lasers, respectively. Images were acquired with following settings: 2048 x 2048 pixels, 16-bit depth. In Figure 4D, fluorescence of emiRFP670 was excited using 640-nm laser. Embryos were imaged by a FV4000 confocal microscope equipped with SilVIR detector (Evident). Images were acquired with following settings: 2048 x 2048 pixels, 16-bit depth.

### Super-resolution imaging with Airyscan2 system

In Figure 1, Figure 2D and H, Figure 4I, Figure S1F and G, data acquisition was performed using Airyscan2 imaging system. Embryos at nc14 were imaged using a Zeiss LSM 900 equipped with Airyscan2 detector. During imaging, temperature was kept in between 22.0 and 23.5°C. Plan-Apochromat 63x / 1.4 N.A. oil immersion objective was used. Images were acquired with following settings: 1180 x 708 pixels (Figure 1, Figure S1G) or 1180 x 732 pixels (Figure 2D and H, Figure 4I) with pixel size 0.043 μm, 16-bit depth, 67 z-slices separated by 0.15 μm. In Figure S1F, GAF-GFP and GAFP-mCherry images were acquired with following settings to achieve optimal Airyscan imaging setup: 1180 x 708 pixels with pixel size 0.043 μm, 16-bit depth, 67 z-slices separated by 0.15 μm for GAF-GFP, 1024 x 612 pixels with pixel size 0.049 μm, 16-bit depth, 56 z-slices separated by 0.18 μm for GAF-mCherry. Fluorescence of mCherry and GFP were excited using 561-nm and 488-nm lasers, respectively. To enable high-speed scanning of embryo samples, Airyscan Multiplex SR-4Y acquisition mode was used. Raw z-stack images were subjected to 3D Airyscan processing using Zen 3.1 software (Zeiss) with the standard mode.

### Scoring of embryo hatching rates

To quantify embryo viability, more than 70 virgin females and males were individually collected for 3 days, mated with each other, and aged for 2 additional days. Following two ∼1-hour pre-lays, embryos were collected for 3 hours and incubated on apple juice plates for ∼48 hours at 25 °C. Subsequently, the number of unhatched embryos was counted to calculate the embryo hatching rate, expressed as the percentage of hatched embryos from total. For embryos laid by *D19B[Δ1]* females, due to the difficulty to visually distinguish unhatched embryos from hatched embryos, the number of hatched embryos was instead measured by counting the number of wandering larvae.

### qRT-PCR

Virgin female flies were collected for 3 days and subsequently fed yeast paste for 3 days before ovary dissection. Ovaries were dissected into 1x PBS buffer. Total RNA was purified from five pairs of ovaries using TRIzol (Thermo Fisher). One microgram of total RNAs was reverse transcribed by PrimeScript RT reagent kit with gDNA eraser (Takara). qRT-PCR was performed using TB Green Premix Ex Taq II (Takara) and the LightCycler 480 System II (Roche). Serial dilutions of cDNA were used for qRT-PCR calibrations. Melting-curve analysis was performed to confirm the amplification of a single product for each target. Following pairs of primers were used for the analysis. *rp49*: (5’-CCG CTT CAA GGG ACA GTA TCT G-3’) and (5’-ATC TCG CCG CAG TAA ACG C-3’). *Burdock*: (5’-GGA CAA ACT CCA TTT CCG ACT C-3’) and (5’-TCC CTG AGC CTG ACT TGT GT-3’). *3S18* (*bel*): (5’-CAA TCG CTT CTT CTC GCT GG-3’) and (5’-CAG GTG TCT GGG GAC TCT TC-3’).

### RNA-seq library preparation

Four hundred ng of total RNA isolated from ovaries was used for rRNA depletion using biotinylated antisense oligos against 18S and 28S rRNAs as previously described.^90^ The rRNA-depleted samples were purified with RNAClean XP beads (Beckman Coulter). Libraries were prepared using the TruSeq Stranded mRNA Library Prep Kit (Illumina, 20020594) according to the manufacturer’s instructions. The multiplexed libraries were sequenced on a NovaSeq X Plus sequencer (Illumina) using paired-end 150 bp reads.

### Computational analysis of RNA-seq data

Adapter sequences were trimmed with Fastp.^69^ The trimmed reads were then aligned to the *Drosophila* melanogaster reference genome (BDGP6.32) using STAR^82^ with the following parameters: --outMultimapperOrder Random --outSAMtype SAM --outReadsUnmapped Fastx --outFilterMultimapNmax 1000 --winAnchorMultimapNmax 1000 --limitOutSAMoneReadBytes 2000000. Sam files were converted to bam files by samtools.^71^ The expression levels of genes and transposable elements were quantified with TEcount^83^ with gene annotation from Ensembl (BDGP6.32) and transposable element annotations downloaded from the TEcount authors’ website (https://www.mghlab.org/software/tetranscripts). Differential expression analysis was performed with DESeq2^84^ for genes and transposable elements with >1 read counts. To quantify the expression of transposon transcripts in *kipf*-depleted versus control ovaries, RNA-seq data from previous publication^14^ was processed following the same protocol.

### Embryo collection and fixation for ChIP

Aged adult flies were discarded from vials, and newly eclosed flies were collected twice: in the evening of the following day and in the morning two days later. Flies were transferred to population cages with apple juice agar plates supplemented with yeast paste and allowed to mate for 1 day at 25 °C. On the following morning, plates were exchanged twice, and embryos were collected over a 30-minute window and aged for 2 hours and 10 minutes at 25 °C. This resulted in embryo collections corresponding to 2 h 10 min-2 h 40 min after egg laying, which corresponds to nc14. Collected embryos were dechorionated by immersion in bleach (Kao) for 2 minutes, followed by thorough rinsing with water. Embryos were fixed by rotating in a mixture of heptane and PBS-T containing 1.8% formaldehyde (final concentration in the aqueous phase) for 10 minutes. Fixation was quenched by adding glycine to a final concentration of 250 mM and rotating for 5 minutes. Embryos were pelleted by centrifugation at 1,000 g for 1 minute at 25 °C. Fixed embryos were flash-frozen in liquid nitrogen and stored at -80 °C until further use.

### Preparation of antibody-conjugated beads

Dynabeads Protein G magnetic beads (Invitrogen) were diluted by adding 10 µL of beads to 100 µL of blocking solution (PBS containing 5 mg/ml BSA). After a brief spin-down, the beads were collected using a magnetic stand and washed twice with blocking solution. The beads were then resuspended in 500 µL of blocking solution, followed by the addition of 2 µL of either anti-FLAG M2 monoclonal antibody (Sigma, F1804) or anti-RNA Polymerase II monoclonal antibody CTD4H8 (Merck Millipore, 05-623). The mixture was rotated at 4 °C for 6 hours. After incubation, the beads were washed three times with blocking solution and finally resuspended in 12 µL of blocking solution.

### ChIP

Approximately 50 µL of nc14 embryos were washed once with PBS-T and centrifuged at 1,000 × *g* for 1 min at 25 °C. The pellet was resuspended in lysis buffer [15 mM HEPES (pH 7.5), 15 mM NaCl, 60 mM KCl, 4 mM MgCl_2_, 0.5% Triton X-100, 0.5 mM DTT, and 1× EDTA-free protease inhibitor (Nacalai Tesque)], and homogenized using a Dounce homogenizer. The lysate was centrifuged at 3,000 *g* for 3 min at 4 °C, and the supernatant was further diluted with lysis buffer and pipetted 30 times to disperse nuclei. The nuclear suspension was sequentially washed once with lysis buffer, wash buffer [15 mM HEPES (pH 7.5), 200 mM NaCl, 1 mM EDTA, 0.5 mM EGTA, and 1× EDTA-free protease inhibitor], and sonication buffer [15 mM HEPES (pH 7.5), 140 mM NaCl, 1 mM EDTA, 0.5 mM EGTA, 0.1% sodium deoxycholate, 0.5% sodium N-lauroylsarcosinate, and 1× EDTA-free protease inhibitor], with 30 pipetting strokes at each step. Nuclei were then resuspended in 300 µL of sonication buffer and subjected to sonication using a Bioruptor Pico (Diagenode) set to 30 sec ON / 30 sec OFF cycles, intensity HIGH, for 7 cycles. Four of the six available holders were used, with standard 1.5 mL tubes. The chromatin fraction was obtained by centrifugation at 20,000 *g* for 15 min at 4 °C. The supernatant was supplemented with 1% Triton X-100 in a total volume of 600 µL. One-twelfth of the sample (50 µL) was reserved as input and stored at -30 °C. Immunoprecipitation was performed overnight at 4 °C by adding the entire volume of antibody-coupled beads (prepared as described above) diluted in 12 µL of blocking solution. Beads were then washed four times with RIPA buffer [50 mM HEPES (pH 7.5), 500 mM LiCl, 1 mM EDTA, 1% NP-40, and 0.7% sodium deoxycholate] and once with TE buffer containing 50 mM NaCl. After centrifugation at 960 *g* for 3 min at 4 °C, the supernatant was completely removed. Beads were resuspended in 200 µL of elution buffer [50 mM Tris-HCl (pH 8.0), 10 mM EDTA, and 1% SDS] and incubated at 65 °C for 30 min. The eluate was collected using a magnetic stand and transferred to a new tube. For the input, 150 µL of elution buffer was added to the 50 µL reserved fraction. De-crosslinking was performed by incubating both ChIP and input samples at 65 °C for 6 hours. Afterward, 200 µL of TE buffer was added. Samples were treated with RNase A (final concentration 10 µg/mL) at 37 °C for 30 min and with proteinase K (final concentration 0.5 mg/mL) at 55 °C for 2 hours to remove RNA and proteins. DNA was purified by phenol-chloroform extraction followed by ethanol precipitation. The final DNA pellet was resuspended in 50 µL of nuclease-free water.

### ChIP-seq library preparation

ChIP-seq libraries were prepared using the NEBNext Ultra II DNA Library Prep Kit for Illumina (New England BioLabs), starting with 15 µL of ChIP DNA and 5 µL of input DNA. During adaptor ligation, the adaptor reagent was diluted 25-fold for ChIP DNA and 10-fold for input DNA. Adapter-ligated DNA was PCR-amplified using Illumina index primers (NEBNext Multiplex Oligos for Illumina, New England BioLabs), with 12 cycles for ChIP DNA and 6 cycles for input DNA. PCR products ranging from 100 to 1,000 bp were size-selected and purified using AMPure XP beads (Beckman Coulter). Size-selected libraries were validated by TapeStation D5000HS kit (Agilent technology). The resulting libraries were sent to BGI Japan for circularization and sequenced on a DNBSEQ-G400 platform (MGI Tech) using paired-end 100 bp reads.

### Computational analysis of ChIP-seq data

Prior to mapping, low-quality reads were removed and adapter sequences were trimmed using Fastp^69^ with the options -q 30 -n 5 -t 1 -l 20 -w 16. The filtered reads were mapped to the *Drosophila melanogaster* reference genome (dm6) using Bowtie2.^70^ Only uniquely mapped reads were retained for analysis using the samtools^71^ view command with the -q 42 option. ChIP-seq experiments were performed with at least two biological replicates. Since the replicates showed high correlation, they were merged and the combined libraries were used for downstream analyses. Peak calling was performed by MACS2^72^ using the input sample as control with -f BAMPE q = 0.01 option, and called peaks were considered as regions of protein binding. Since ChIP-seq of insulator proteins such as CTCF are known to tend to give rise to false-positive peaks, often referred to as “phantom peaks,”^91^ ChIP-seq using anti-FLAG M2 monoclonal antibody (Sigma, F1804) for the control *yw* strain (the parental strain for all GFP-3xFLAG tag knock-in strains) was performed to identify such peaks. In total, 269 genomic regions were identified as phantom peaks (summarized in Table S2). Any peak in the experimental samples that overlapped by even 1 bp with these phantom peaks was excluded from further analysis. Overlaps were determined using the bedtools intersect command.^73^ For heatmaps and profile plots, ChIP scores were calculated using deepTools.^74^ BigWig files were generated from BAM files using bamCoverage or bamCompare, and matrix data were computed using computeMatrix, followed by plotting. In bamCompare, input samples were used as control for signal subtraction. All genome browser views and heatmap visualizations used ChIP tracks with input-subtracted signals. Correlation of mapping files were confirmed using the multiBamSummary program in deepTools. For upset plots, the narrowPeak files obtained from MACS2 callpeak were processed using the Granges function in the GenomicRanges package,^75^ and visualized with the UpSetR package.^76^ Quantitative comparison of ChIP peaks was performed using MAnorm.^77^ For visualization of ChIP-seq tracks, Integrative Genome Viewer^78^ was used. For visualization of ChIP-seq tracks alongside Micro-C data, CoolBox^66^ was used. For Figure 7D, the file GSE171396_Batut_MicroC_yw_CC14_Boundaries_min1.25.bed.gz from the GEO dataset GSE171396 was utilized to define the genomic positions and strengths of topological boundaries.^50^ ChIP-seq quality metrics, including alignment rates, were summarized in Table S2. As a quality metrics for ChIP-seq, library complexity (the proportion of unique reads in the entire library, indicating low levels of PCR bias) was calculated using bam files by the DROMPA3^79^ software. To evaluate the signal-to-noise ratio of the ChIP signal, the strand-shift profile and synthetic Jensen-Shannon Divergence were calculated using bam files by SSP^80^ and plotFingerprint program in deepTools, respectively.

### Analysis of zinc-finger domain similarity

Domain compositions were defined following the annotations provided in the UniProt database for each protein. To assess the similarity of C2H2 zinc-finger domains among D19A, D19B and BroN, C2H2 motifs were further identified using the regular expression “(.{2})C(.{2}|.{4})C(.{12})H(.{3,5})H” (reviewed in Pabo et al. ^92^) with custom Python code. Each zinc-finger domain in D19B or BroN was then aligned to the corresponding zinc-finger domain in D19A using BioPython^81^ ensuring that core Cys and His residues were always aligned with each other. The alignment results are presented in Table S3. Similarity scores were defined as the percentage of perfectly aligned residues in the zinc-finger domain of interest, relative to the total number of residues in the corresponding D19A zinc-finger domain.

### Computational analysis of Micro-C data

Micro-C from a previous publication^50^ was analyzed using the 4DN Hi-C analysis pipeline (https://data.4dnucleome.org/resources/data-analysis/hi_c-processing-pipeline). In brief, adapter sequences were trimmed from paired-end reads with Fastp,^69^ and then reads were mapped to the *Drosophila melanogaster* reference genome (BDGP6.32) using BWA^85^ with the option -SP5M. Alignments were filtered to retain reads with alignment quality ≥3, sorted, and deduplicated using pairtools.^86^ The deduplicated pairs were then selected using pairtools with the query ‘(pair_type == “UU”) or (pair_type == “UR”) or (pair_type == “RU”)’. Matrix aggregation was performed with Cooler.^87^

## Plasmids

### pBlueScript-CG30020^WT^ CDS

A DNA fragment containing *CG30020* CDS was PCR amplified from *yw* cDNA using primers (5’-TTT AAG TCG ACA TGG CGC TGA ACA AAA GGA AGA GC-3’) and (5’-TTA AAG AAT TCG GGC TGC CCA TTT TCT TCG C-3’) and inserted between the SalI and EcoRI sites of pBlueScript SK II (-) plasmid using corresponding restriction enzymes.

### pBlueScript-CG30020^ΔZAD^ CDS

Coding sequence for ZAD domain was specifically removed from pBlueScript-CG30020^WT^ CDS plasmid by PCR-based site-directed mutagenesis using primers (5’-TTC GGG CGC CGA AGT GGA CTA CAA GCT ATG TCC CAA GGT TTC AAA GAG AC-3’) and (5’-GTC TCT TTG AAA CCT TGG GAC ATA GCT TGT AGT CCA CTT CGG CGC CCG AA-3’).

### pBlueScript-Nnk^WT^ CDS

DNA fragment containing *Nnk* CDS was PCR amplified from *yw* cDNA using primers (5’-AAA TTC TCG AGA TGA CGC CAC AAT GCC GGT TG-3’) and (5’-AAT TTC TGC AGT TTA TTT TTG GAA TCA ATT TCG G-3’) and inserted between the XhoI and PstI sites of pBlueScript SK II (-) plasmid using corresponding restriction enzymes.

### pBlueScript-Nnk^ΔZAD^ CDS

Coding sequence for ZAD domain was specifically removed from pBlueScript-Nnk^WT^ CDS plasmid by PCR-based site-directed mutagenesis using primers (5’-TAC CGG GCC CCC CCT CGA GAT GAC GCG CCG AAG TGT GGA AGC CAA GGA TG-3’) and (5’-CAT CCT TGG CTT CCA CAC TTC GGC GCG TCA TCT CGA GGG GGG GCC CGG TA-3’).

### pBlueScript-D19B^WT^ CDS

A DNA fragment containing *D19B* CDS was PCR amplified from *yw* cDNA using primers (5’-AAA TTG TCG ACA TGA ATG AGG AGA GCC AGT ACT CC-3’) and (5’-AAT TTG AAT TCA ACA TGA TCC AAC TTC AGT GCG TC-3’) and inserted between the SalI and EcoRI sites of pBlueScript SK II (-) plasmid using corresponding restriction enzymes.

### pBlueScript-D19B^ΔZAD^ CDS

Coding sequence for ZAD domain was specifically removed from pBlueScript-D19B^WT^ CDS plasmid by PCR-based site-directed mutagenesis using primers (5’-CAG TAC TCC ATA CAC GTG GCC AGC AAC GCG-3’) and (5’-CGC GTT GCT GGC CAC GTG TAT GGA GTA CTG-3’).

### pBlueScript-5’region-CG30020^WT^ CDS-GFP-3xFLAG-3’region

A DNA fragment containing ∼1.5-kb 3’ regulatory region (including 3’ UTR and polyA signal) of *CG30020* was PCR amplified from *yw* genomic DNA and inserted between the EcoRI and NotI sites of pBlueScript-CG30020^WT^ CDS. Subsequently, a DNA fragment containing GFP-3xFLAG was PCR amplified and inserted between the EcoRI and BglII sites. Lastly, a DNA fragment containing ∼2.1-kb 5’ regulatory region of *CG30020* (including TSS and 5’ UTR) was PCR amplified from *yw* genomic DNA and inserted between the KpnI and SalI sites.

### pBlueScript-5’region-CG30020^ΔZAD^ CDS-GFP-3xFLAG-3’region

pBlueScript-CG30020^ΔZAD^ CDS was digested with SalI and EcoRI, and resulting DNA fragment containing CG30020^ΔZAD^ CDS was inserted between the SalI and EcoRI sites of pBlueScript-5’region-CG30020^WT^ CDS-GFP-3xFLAG-3’region plasmid.

### pbφ-CG30020^WT^ CDS-GFP-3xFLAG

pBlueScript-5’region-CG30020^WT^ CDS-GFP-3xFLAG-3’region plasmid was digested with SpeI and NotI, and resulting DNA fragment was inserted into the pbφ vector.

### pbφ-CG30020^ΔZAD^ CDS-GFP-3xFLAG

pBlueScript-5’region-CG30020^ΔZAD^ CDS-GFP-3xFLAG-3’region plasmid was digested with SpeI and NotI, and resulting DNA fragment was inserted into the pbφ vector.

### pBlueScript-5’region-Nnk^WT^ CDS-GFP-3xFLAG-3’region

A DNA fragment containing ∼1.4-kb 3’ regulatory region (including 3’ UTR and polyA signal) of *Nnk* was PCR amplified from *yw* genomic DNA and inserted between the PstI and NotI sites of pBlueScript-Nnk^WT^ CDS. Subsequently, a DNA fragment containing GFP-3xFLAG was PCR amplified and inserted between the PstI and BglII sites. Lastly, a DNA fragment containing ∼1.9-kb 5’ regulatory region of *Nnk* (including TSS and 5’ UTR) was PCR amplified from *yw* genomic DNA and inserted between the ApaI and XhoI sites.

### pBlueScript-5’region-Nnk^ΔZAD^ CDS-GFP-3xFLAG-3’region

pBlueScript-Nnk^ΔZAD^ CDS was digested with XhoI and PstI, and resulting DNA fragment containing Nnk^ΔZAD^ CDS was inserted between the XhoI and PstI sites of pBlueScript-5’region-Nnk^WT^ CDS-GFP-3xFLAG-3’region plasmid.

### pbφ-Nnk^WT^ CDS-GFP-3xFLAG

pBlueScript-5’region-Nnk^WT^ CDS-GFP-3xFLAG-3’region plasmid was digested with SpeI and NotI, and resulting DNA fragment was inserted into the pbφ vector.

### pbφ-Nnk^ΔZAD^ CDS-GFP-3xFLAG

pBlueScript-5’region-Nnk^ΔZAD^ CDS-GFP-3xFLAG-3’region plasmid was digested with SpeI and NotI, and resulting DNA fragment was inserted into the pbφ vector.

### pBlueScript-attB-D19B CDS^WT^-GFP-3xFLAG-attB

A DNA fragment containing ∼800-bp 5’ regulatory region of *D19B* (including TSS and 5’ UTR) was PCR amplified from *yw* genomic DNA and inserted between the ApaI and XhoI sites of pBlueScript-attB-MCS-GFP-3xFLAG-attB plasmid. Subsequently, a DNA fragment containing ∼700-bp 3’ regulatory region (including 3’ UTR and polyA signal) of *D19B* was PCR amplified from *yw* genomic DNA and inserted between the XbaI and NotI sites of the plasmid. Lastly, a DNA fragment containing *D19B^WT^* CDS was PCR amplified from pBlueScript-D19B^WT^ CDS and inserted between the SalI and EcoRI sites of the plasmid.

### pBlueScript-attB-D19B CDS^ΔZAD^-GFP-3xFLAG-attB

A DNA fragment containing ∼800-bp 5’ regulatory region of *D19B* (including TSS and 5’ UTR) was PCR amplified from *yw* genomic DNA and inserted between the ApaI and XhoI sites of pBlueScript-attB-MCS-GFP-3xFLAG-attB plasmid. Subsequently, a DNA fragment containing ∼700-bp 3’ regulatory region (including 3’ UTR and polyA signal) of *D19B* was PCR amplified from *yw* genomic DNA and inserted between the XbaI and NotI sites of the plasmid. Lastly, a DNA fragment containing *D19B^ΔZAD^* CDS was PCR amplified from pBlueScript-D19B^ΔZAD^ CDS and inserted between the SalI and EcoRI sites of the plasmid.

### pbφ-His2Av-emiRFP670

A DNA fragment containing emiRFP670 was PCR amplified from pH2B-emiRFP670 (addgene# 136571) and inserted between the HindIII and NheI sites of pbφ-His2Av-eBFP2 plasmid.^93^

## Supplemental Tables

**Table S1. DNA oligos used to build gRNA expression plasmids, related to Figure 1, S1, S2.**

**Table S2. List of false-positive anti-FLAG ChIP-seq peaks and ChIP-seq quality metrics, related to Figure 3, 5, 6, 7, S2, S3, S4, S6, S7.**

**Table S3. Analysis of zinc-finger domain similarity, related to Figure 5.**

