## Supplemental Figure for "Decoding the molecular logic of rapidly evolving ZAD zinc-finger proteins in *Drosophila*"

### Figure S1

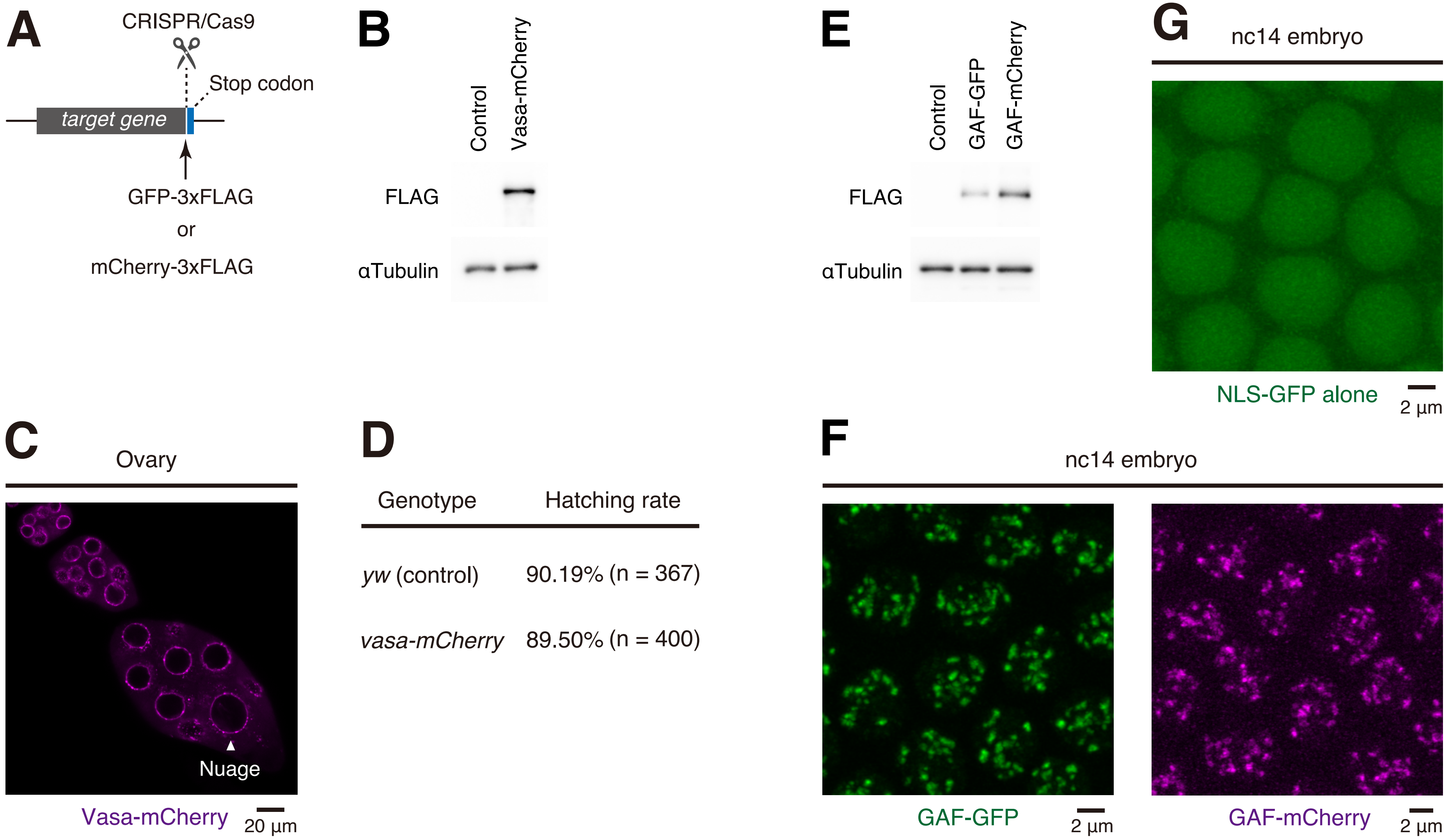
A
CRISPR/Cas9
Stop codon
target gene
GFP-3xFLAG
or
mCherry-3xFLAG
B
Control
Vasa-mCherry
FLAG
αTubulin
E
Control
GAF-GFP
GAF-mCherry
FLAG
αTubulin
G
nc14 embryo
NLS-GFP alone
2 µm
C
Ovary
Nuage
Vasa-mCherry
20 µm
D
Genotype
Hatching rate
*yw* (control)
90.19% (n = 367)
*vasa-mCherry*
89.50% (n = 400)
F
nc14 embryo
GAF-GFP
2 µm
GAF-mCherry
2 µm

**Figure S1. Establishment of CRISPR/Cas9-mediated protein-tagging system, Related to Figure 1.**

(A) Schematic representation of CRISPR/Cas9-mediated protein-tagging system.

(B) Western blot analysis of mCherry-3xFLAG-tagged endogenous Vasa. Two-to-four hour embryos from homozygous *vasa-mCherry-3xFLAG* genome-edited strain were used for the analysis. As a control, 2-4 h *yw* embryos were used.

(C) Confocal imaging of Vasa-mCherry-3xFLAG in dissected ovary. A single z-plane image is shown.

(D) Measurement of the embryo hatching rate. Embryos from homozygous *vasa-mCherry-3xFLAG* genome-edited strain were used for the analysis.

(E) Western blot analysis of GFP-3xFLAG- or mCherry-3xFLAG-tagged endogenous GAF. Two-to-four hour embryos from corresponding homozygous genome-edited strain were used for the analysis. As a control, 2-4 h *yw* embryos were used.

(F) Airyscan imaging of GFP-3xFLAG- or mCherry-3xFLAG-tagged endogenous GAF in living embryos. Images were taken ~15 min after entry into nc14. Embryos from corresponding homozygous genome-edited strains were used for the analysis. The maximum intensity projected images are shown.

(G) Airyscan imaging of NLS-GFP-3xFLAG fusion protein in living embryos. Images were taken ~15 min after entry into nc14. Embryos from homozygous *NLS-GFP-3xFLAG* transgenic strain were used for the analysis. The maximum intensity projected images are shown.

Figure S2

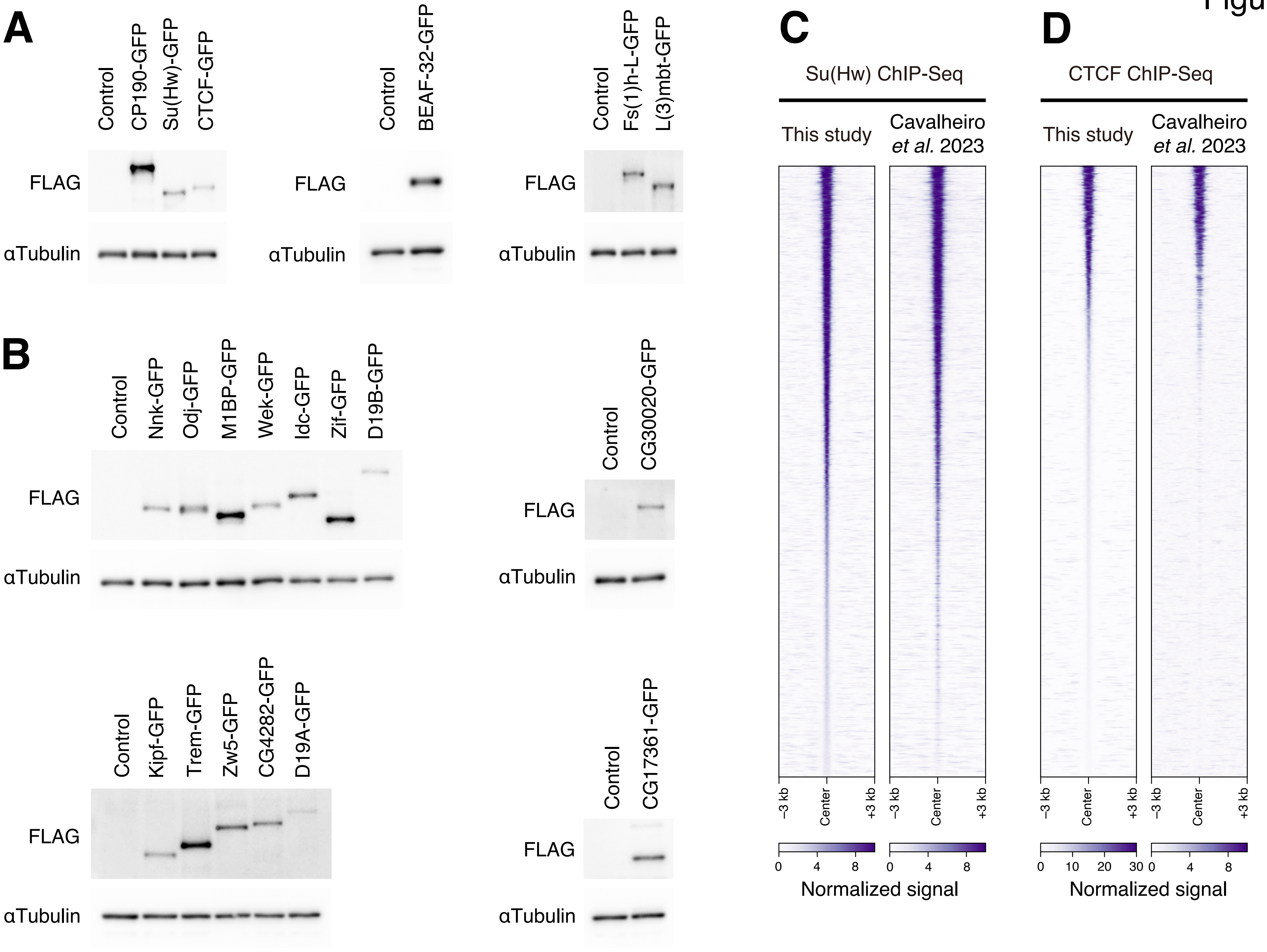
A
Control
CP190-GFP
Su(Hw)-GFP
CTCF-GFP
FLAG
αTubulin
Control
BEAF-32-GFP
FLAG
αTubulin
Control
Fs(1)h-L-GFP
L(3)mbt-GFP
FLAG
αTubulin
B
Control
Nnk-GFP
Odj-GFP
M1BP-GFP
Wek-GFP
Idc-GFP
Zif-GFP
D19B-GFP
FLAG
αTubulin
Control
CG30020-GFP
FLAG
αTubulin
Control
Kipf-GFP
Trem-GFP
Zw5-GFP
CG4282-GFP
D19A-GFP
FLAG
αTubulin
Control
CG17361-GFP
FLAG
αTubulin
C
Su(Hw) ChIP-Seq
This study
Cavalheiro et al. 2023
−3 kb
Center
+3 kb
0
4
8
Normalized signal
D
CTCF ChIP-Seq
This study
Cavalheiro et al. 2023
−3 kb
Center
+3 kb
0
10
20
30
4
8
Normalized signal

**Figure S2. Tagging of endogenous insulator proteins and ZAD-ZnFs, Related to Figure 1.**

(A) Western blot analysis of GFP-3xFLAG-tagged endogenous insulator proteins. Two-to-four hour embryos from corresponding homozygous genome-edited strains were used for the analysis. As a control, 2-4 h *yw* embryos were used.

(B) Western blot analysis of GFP-3xFLAG-tagged endogenous ZAD-ZnFs. Two-to-four hour embryos from corresponding homozygous genome-edited strains were used for the analysis. As a control, 2-4 h *yw* embryos were used.

(C) Heatmap visualization of Su(Hw) ChIP-seq peaks detected in our ChIP-seq analysis using anti-FLAG antibody (left) and previously reported ChIP-seq analysis using anti-Su(Hw) antibody (right).[43]

(D) Heatmap visualization of CTCF ChIP-seq peaks detected in our ChIP-seq analysis using anti-FLAG antibody (left) and previously reported ChIP-seq analysis using anti-CTCF antibody (right).[43]

Figure S3

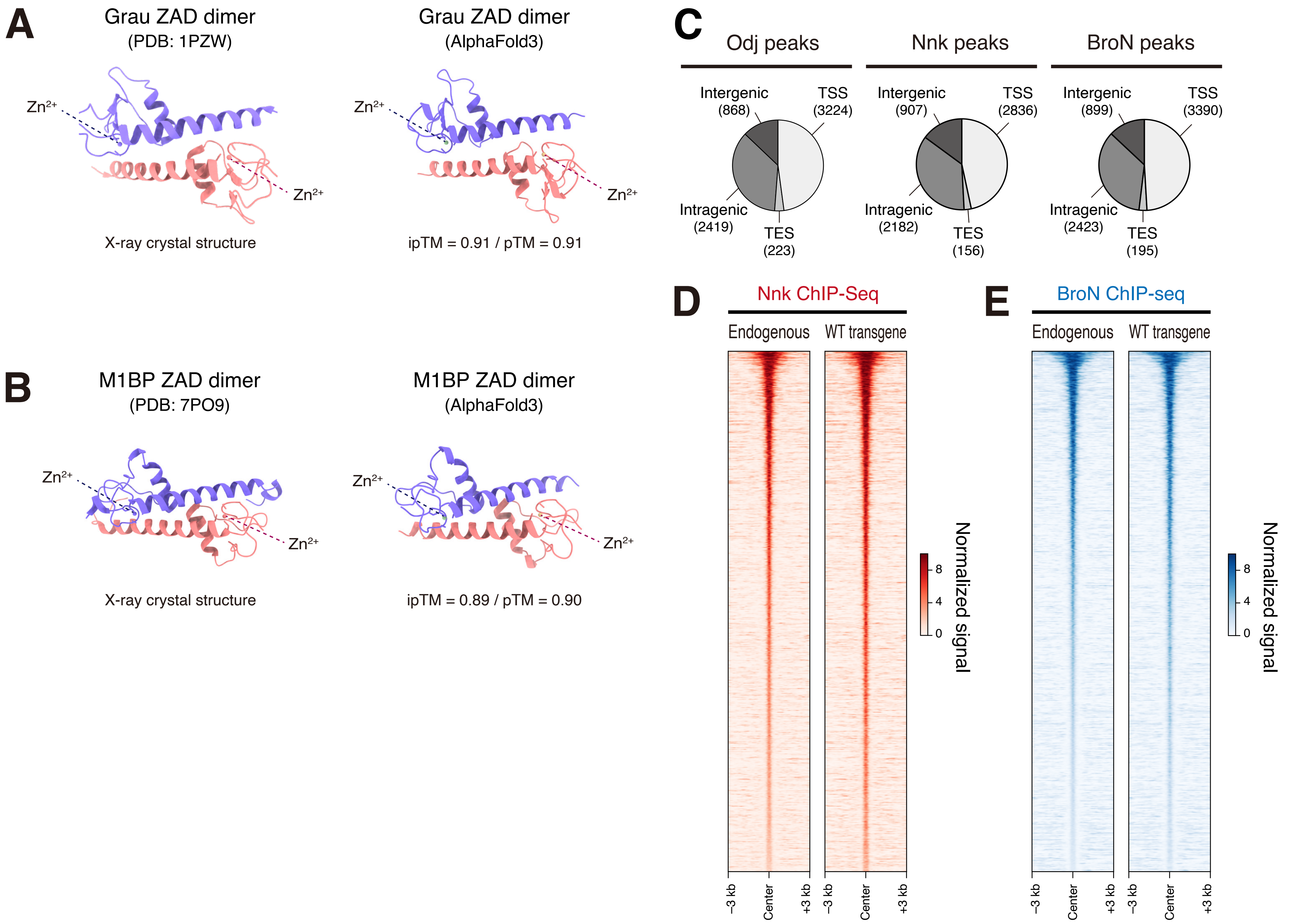
A
Grau ZAD dimer
(PDB: 1PZW)
Zn2+
Zn2+
X-ray crystal structure
Grau ZAD dimer
(AlphaFold3)
Zn2+
Zn2+
ipTM = 0.91 / pTM = 0.91
B
M1BP ZAD dimer
(PDB: 7PO9)
Zn2+
Zn2+
X-ray crystal structure
M1BP ZAD dimer
(AlphaFold3)
Zn2+
Zn2+
ipTM = 0.89 / pTM = 0.90
C
Odj peaks
Intergenic (868)
TSS (3224)
Intragenic (2419)
TES (223)
Nnk peaks
Intergenic (907)
TSS (2836)
Intragenic (2182)
TES (156)
BroN peaks
Intergenic (899)
TSS (3390)
Intragenic (2423)
TES (195)
D
Nnk ChIP-Seq
Endogenous
WT transgene
−3 kb
Center
+3 kb
−3 kb
Center
+3 kb
8
4
0
Normalized signal
E
BroN ChIP-seq
Endogenous
WT transgene
−3 kb
Center
+3 kb
−3 kb
Center
+3 kb
8
4
0
Normalized signal

**Figure S3. AlphaFold3 prediction of ZAD dimers, Related to Figure 2 and 3.**

(A) X-ray crystal structure (left, PDB: 1PZW) and AlphaFold3 prediction (right) of the Grau ZAD dimer. Protein structure was visualized using UCSF ChimeraX.[65]

(B) X-ray crystal structure (left, PDB: 7PO9) and AlphaFold3 prediction (right) of the M1BP ZAD dimer. Protein structure was visualized using UCSF ChimeraX.[65]

(C) Pie chart showing the distribution profiles of endogenous Odj, Nnk, and BroN ChIP-seq peaks.

(D) Heatmap visualization of ChIP-seq peaks endogenous Nnk (left) and WT Nnk transgene product (right)

(E) Heatmap visualization of ChIP-seq peaks of endogenous BroN (left) and WT BroN transgene product (right).

Figure S4

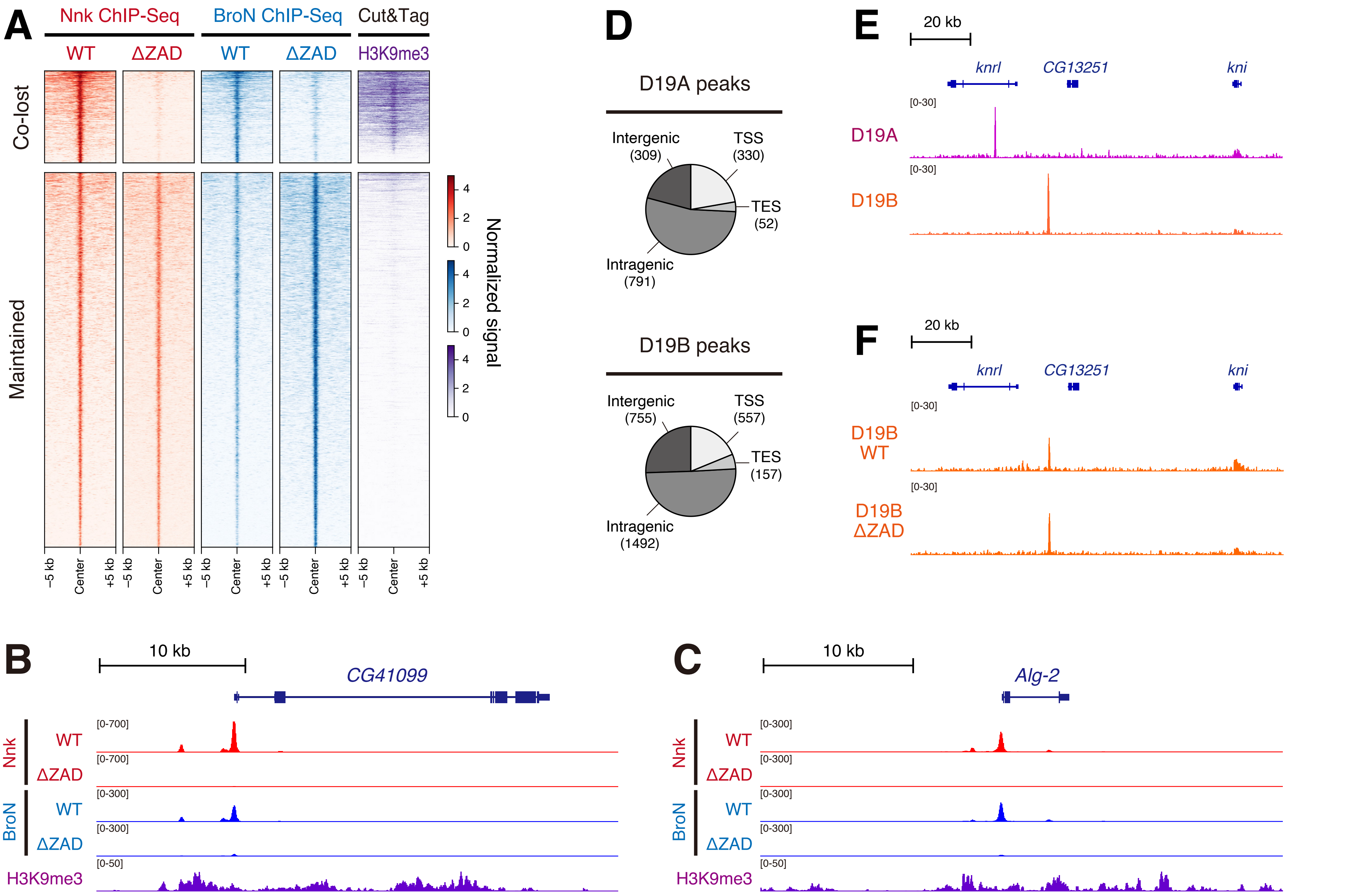
A
Nnk ChIP-Seq
BroN ChIP-Seq
Cut&Tag
WT
ΔZAD
WT
ΔZAD
H3K9me3
Co-lost
Maintained
Normalized signal
–5 kb
Center
+5 kb
D
D19A peaks
Intergenic (309)
TSS (330)
TES (52)
Intragenic (791)
D19B peaks
Intergenic (755)
TSS (557)
TES (157)
Intragenic (1492)
E
20 kb
knrl
CG13251
kni
[0-30]
D19A
D19B
F
20 kb
D19B WT
D19B ΔZAD
B
10 kb
CG41099
[0-700]
[0-300]
[0-50]
Nnk
BroN
H3K9me3
C
10 kb
Alg-2

**Figure S4. ZAD-dependent recruitment of Nnk and BroN to H3K9me3-enriched heterochromatic regions, Related to Figure 3 and 4.**

(A) Heatmap visualization of WT and ΔZAD Nnk/BroN ChIP-seq peaks along with H3K9me3 Cut&Tag signal[48] sorted by the strength of WT Nnk.

(B and C) IGV genome browser tracks showing WT/ΔZAD Nnk and WT/ΔZAD BroN ChIP-seq coverage at the H3K9me3-enriched region.

(D) Pie chart showing distribution profiles of endogenous D19A and D19B ChIP-seq peaks.

(E) IGV genome browser tracks showing endogenous D19A and D19B ChIP-seq coverage.

(F) IGV genome browser tracks showing WT and ΔZAD D19B ChIP-seq coverage.

### Figure S5

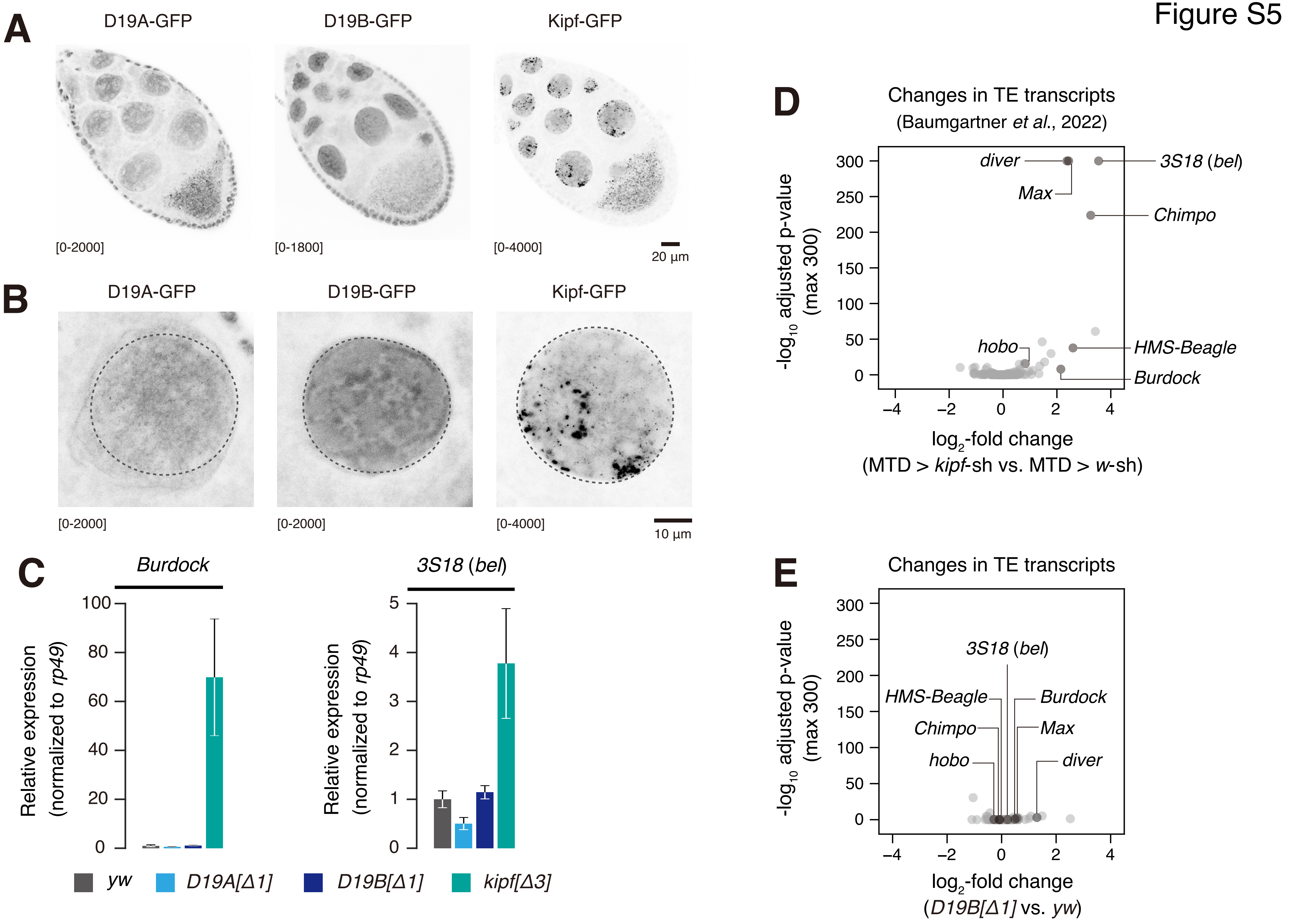

A
D19A-GFP
D19B-GFP
Kipf-GFP
[0-2000]
[0-1800]
[0-4000]
20 µm
B
D19A-GFP
D19B-GFP
Kipf-GFP
[0-2000]
[0-2000]
[0-4000]
10 µm
C
Burdock
3S18 (bel)
Relative expression (normalized to rp49)
100
80
60
40
20
0
5
4
3
2
1
0
yw
D19A[Δ1]
D19B[Δ1]
kipf[Δ3]
D
Changes in TE transcripts
(Baumgartner et al., 2022)
-log10 adjusted p-value (max 300)
300
250
200
150
100
50
0
diver
Max
3S18 (bel)
Chimpo
hobo
HMS-Beagle
Burdock
−4
−2
0
2
4
log2-fold change
(MTD > kipf-sh vs. MTD > w-sh)
E
Changes in TE transcripts
-log10 adjusted p-value (max 300)
3S18 (bel)
HMS-Beagle
Burdock
Chimpo
Max
hobo
diver
log2-fold change
(D19B[Δ1] vs. yw)

**Figure S5. Loss of D19A and D19B does not lead to derepression of transposable elements, Related to Figure 4 and 5.**

(A) Confocal imaging of endogenous D19A-, D19B-, and Kipf-GFP-3xFLAG in dissected ovary. The maximum intensity projection of thin sliced z-stack images is shown.

(B) High-magnification view of endogenous D19A-, D19B-, and Kipf-GFP-3xFLAG in nurse cell nucleus. The maximum intensity projection of thin sliced z-stack images is shown. The same original imaging data shown in (A) were used for the analysis.

(C) The expression levels of *Burdock* and *3S18* (*bel*) transposons in dissected ovary of homozygous *D19A[Δ1]*, *D19B[Δ1]*, *kipf[Δ3]* and control *yw* virgin female. Expression levels were normalized to *rp49* mRNA. Error bars represent mean ± standard deviation of three biological replicates.

(D) Volcano plot depicting the $\log_2$-fold changes in transposon transcripts in *kipf*-depleted versus control ovaries. Publicly available RNA-seq data (Baumgartner *et al.*[14] ; SRR19139212, SRR19139213, SRR19139214, SRR19139218, SRR19139219, SRR19139220) was used for the analysis.

(E) Volcano plot depicting the $\log_2$-fold changes in transposon transcripts in homozygous *D19A[Δ1]* versus control *yw* ovaries.

### Figure S6

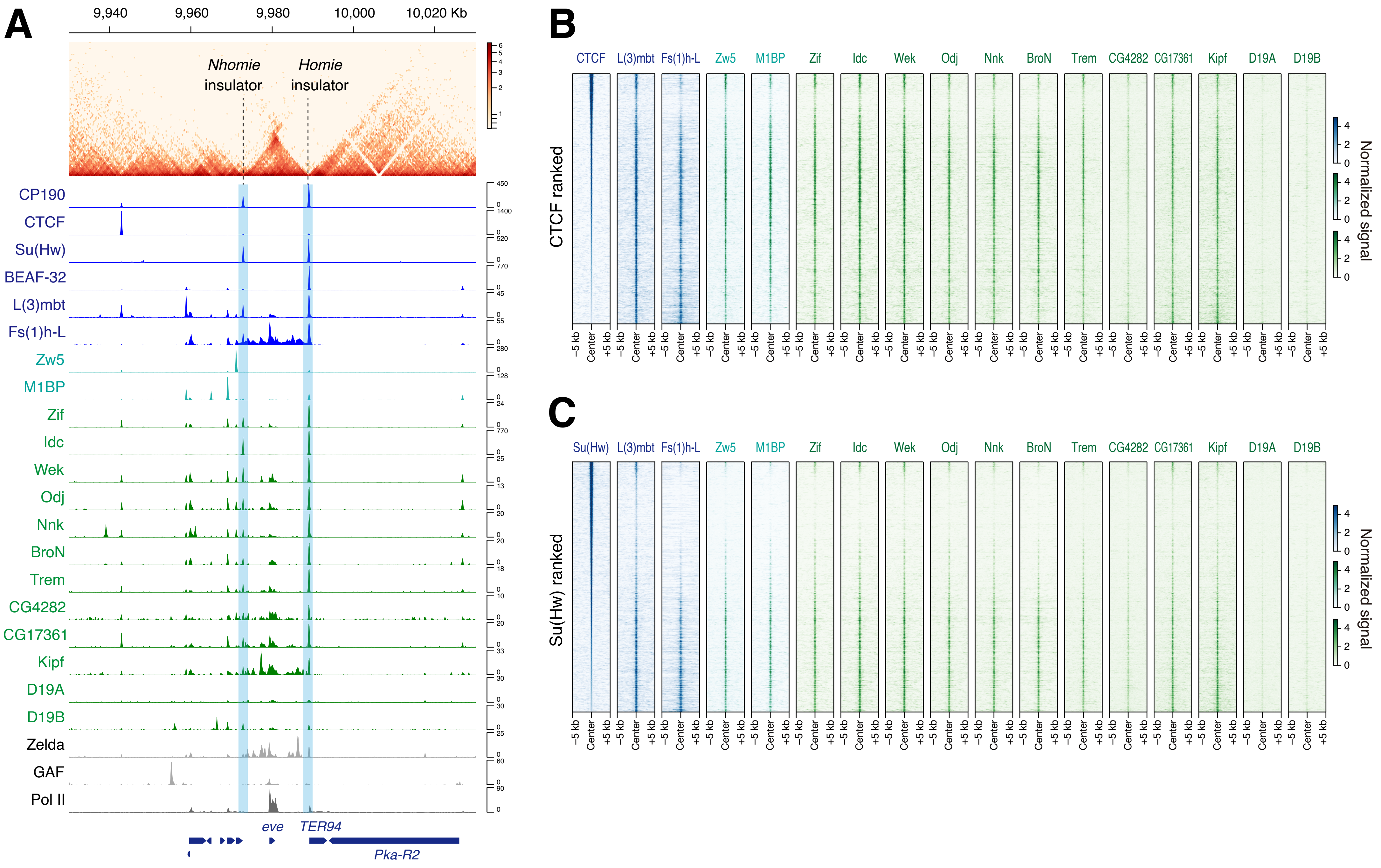

A
9,940
9,960
9,980
10,000
10,020 Kb
Nhomie insulator
Homie insulator
CP190
CTCF
Su(Hw)
BEAF-32
L(3)mbt
Fs(1)h-L
Zw5
M1BP
Zif
Idc
Wek
Odj
Nnk
BroN
Trem
CG4282
CG17361
Kipf
D19A
D19B
Zelda
GAF
Pol II
eve
TER94
Pka-R2
B
CTCF ranked
CTCF
L(3)mbt
Fs(1)h-L
Zw5
M1BP
Zif
Idc
Wek
Odj
Nnk
BroN
Trem
CG4282
CG17361
Kipf
D19A
D19B
−5 kb
Center
+5 kb
Normalized signal
C
Su(Hw) ranked
Su(Hw)

**Figure S6. ZAD-ZnFs possess intrinsic insulator binding activity, Related to Figure 6 and 7.**

(A) Coolbox toolkit[66] was used to visualize Micro-C map along with ChIP-seq profiles of ZAD-ZnFs and non-ZAD-ZnF insulator proteins at the *eve* locus. Publicly available Micro-C data in nc14 embryos[50] was used for the analysis.

(B) Heatmap visualization of ChIP-seq profiles of non-ZAD-ZnF insulator proteins and ZAD-ZnFs sorted by the strength of CTCF ChIP-seq peaks.

(C) Heatmap visualization of ChIP-seq profiles of non-ZAD-ZnF insulator proteins and ZAD-ZnFs sorted by the strength of Su(Hw) ChIP-seq peaks.

Figure S7

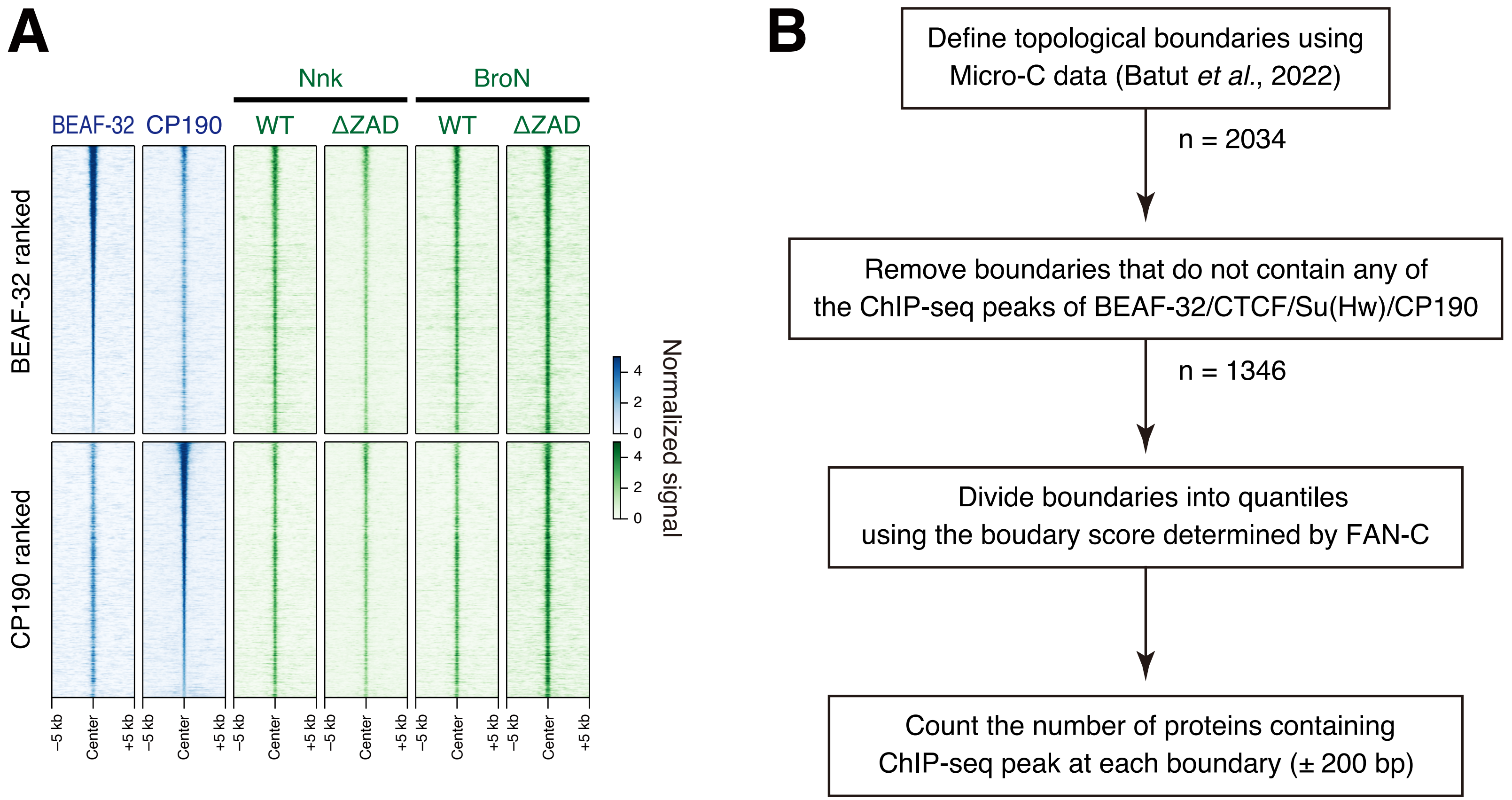


List of 20 proteins used for the analysis:

**BEAF-32, CTCF, Su(Hw), CP190, L(3)mbt, Fs(1)h-L, Zw5, M1BP, Zif Idc, Wek, Odj, Nnk, BroN, Trem, CG4282, CG17361, Kipf, D19A, D19B**

*False-positive peaks are initially removed using the control *yw* ChIP-seq data for each dataset

**Figure S7. Nnk and BroN associate with insulator elements independently of ZAD, Related to Figure 7.**

(A) Heatmap visualization of BEAF-32, CTCF, WT/ΔZAD Nnk and WT/ΔZAD BroN ChIP-seq profiles sorted by the strength of BEAF-32 (upper panels) or CTCF (lower panels) ChIP-seq peaks.

(B) Schematic representation of the analysis shown in Figure 7D.
